# Repeated extrinsic and anisotropic mechanical inputs promote polarized adherens junction elongation

**DOI:** 10.1101/2024.07.09.602689

**Authors:** Xinyi Yang, Teresa Ferraro, Kelly Molnar, Julien Pontabry, Sam-Rayden Malanda, Nicola Maghelli, Loïc Royer, Stephan W. Grill, Gene Myers, Silvia Grigolon, Michel Labouesse

## Abstract

A key challenge in development is understanding how complex organisms physically coordinate the morphogenesis of multiple tissues. Here, using biophysical approaches, we investigate how muscles under the epidermis specifically stimulate the extension of anterior-posterior (AP-oriented) epidermal adherens junctions during late *C. elegans* embryonic elongation. First, light-sheet imaging shows that asynchronous patterns of muscle contractions drive embryo rotations. In turn, junctions between the lateral and dorso-ventral epidermis repeatedly oscillate between a folded, hypotensed state, and an extended, hypertensed state. Second, FRAP (Fluorescence Recovery After Photobleaching) analysis of an E-cadherin::GFP construct shows that muscle contractions stimulate E-cadherin turnover. Moreover, a mechano-chemical model backed by genetic tests suggests that E-cadherin trafficking modulates junction elongation due to lower line tension. Altogether, our results illustrate how muscle contractions fluidize epidermal adherens junctions, which, combined with anisotropic tension in the epidermis, drives their polarized extension.

## INTRODUCTION

Morphological changes in one tissue can mechanically affect nearby tissues through pressure or tensile stress (Goodwin and Nelson, 2021; Julicher and Eaton, 2017; Stooke-Vaughan and Campas, 2018). For instance, these mechanical stimuli can impact fundamental cellular processes such as cell division orientation, cytoskeleton dynamics, junction turnover, or planar polarity (Aigouy et al., 2010; Levayer et al., 2011; Pare et al., 2019; Rauzi et al., 2010; van Leen et al., 2020). While we appreciate their importance, the nature of such mechanical feedback often requires further definition.

*C. elegans* embryos offer a powerful system for studying the influence of multiple tissues on morphogenesis. Its external epithelium comprises two main cell types, lateral epidermal cells enriched in myosin II, and adjacent dorsal and ventral epidermal cells forming circumferentially oriented actin bundles. The latter are tightly mechanically coupled to four longitudinally oriented muscle rows via hemidesmosomes (Zhang and Labouesse, 2010) (Fig. 1A-A’; Fig. S1A-A”). Embryonic elongation includes two phases: until the so-called 2-fold stage, elongation depends on tension and stiffness anisotropies in the epidermis (Vuong-Brender et al., 2017); beyond, it also requires muscles (Williams and Waterston, 1994) (Fig. 1A-A’). During the second stage, local muscle contractions elicit mechanotransduction signaling in the epidermis (Zhang et al., 2011), promoting hemidesmosome maturation (Zahreddine et al., 2010; Zhang et al., 2011) and triggering actin reorganization (Fig. 1A’’) (Gillard et al., 2019; Lardennois et al., 2019). In muscle contraction-defective mutants (called Pat mutants), these events are disrupted (Gillard et al., 2019; Lardennois et al., 2019; Zahreddine et al., 2010; Zhang et al., 2011), and embryos stop elongating at the 2-fold stage (Williams and Waterston, 1994).

**Figure 1.**
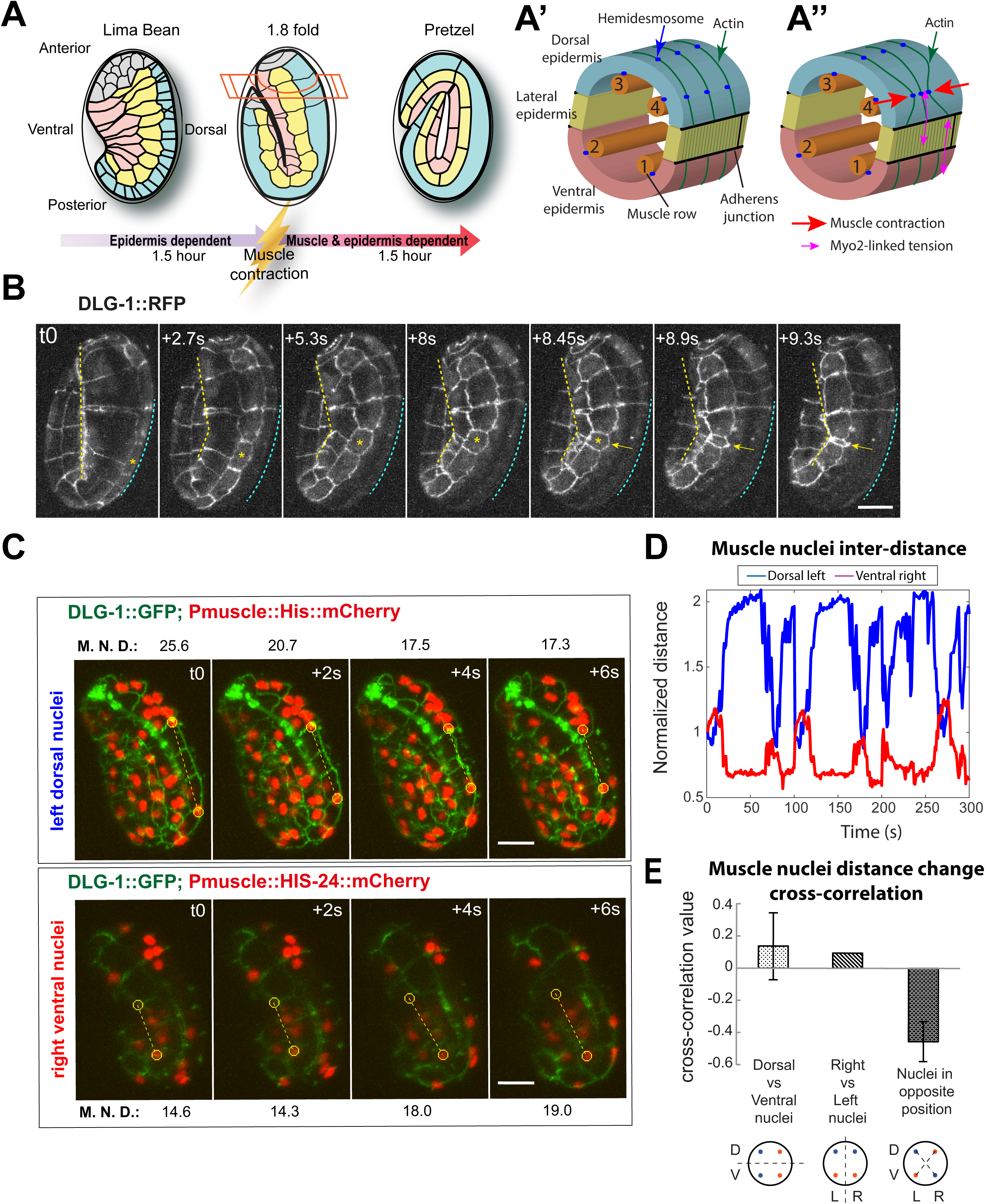
Alternate muscle activity breaks the symmetry of embryo. **(A-A’’)** Scheme of embryo elongation and junction remodeling (black lines in A), and cross-section through the embryo at the level of the head in the 1.8-fold embryo (A’; internal organs omitted for clarity). The muscle-independent and -dependent phases are ≈1.5 hour each. The 81 embryonic muscle cells are divided into four areas (1-4); hemidesmosomes (blue dots, see Fig. S1A-A’’) tightly link the epidermis to muscles, such that muscles locally deflect the actin cytoskeleton when contracting (red arrows in A’’), see (Lardennois et al., 2019). **(B)** Still pictures from a lightsheet movie of the marker DLG-1::RFP (strain ML1652) illustrating embryo rotation within the eggshell (see Movie S1): the V1 lateral cell (yellow asterisk) travels from the outer edge (cyan dotted line) to the fold (yellow dotted line), which we call inward rotation (in outward rotations, lateral cells travel to the outer edge). Timing in upper left corner. **(C)** Still pictures from a two-color lightsheet movie (strain ML2616) illustrating how contralateral muscles contract alternatively (Movie S2); upper and lower panels, projections showing the left and right sides, respectively. Numbers above or below the images give the distance in µm between the circled muscle nuclei (M.N.D., see yellow dotted line). **(D)** Normalized distance change between two selected muscle nuclei on dorsal left (blue) and ventral right muscles (red). **(E)** Muscle nuclei distance cross-correlation between dorsal and ventral, left and right and contralateral nuclei (left-ventral and right-dorsal or right-ventral and life-dorsal sides) of the embryo (N=4 embryos). Scale bar, 10 µm. All embryos oriented with anterior up. See also Fig. S1.

Here we address two inter-related questions. One is to explain how muscle activity also specifically promotes the extension of anterior-posterior (AP) oriented adherens junctions, which is blocked in Pat mutants. Another is to determine the link between muscle activity and junction turnover. To do so, we combine several microscopy modes to get a global view of junction dynamics. We further develop a systematic kinetic analysis of E-cadherin turnover, together with a physical model to explain how muscles promote the preferential extension of AP-oriented junctions by facilitating E-cadherin exocytosis and reducing line tension. By comparing our data with data from other organisms, we suggest some general features for the influence of repeated mechanical inputs on adherens junctions.

## RESULTS

### Muscle contractions transiently change epidermal cell shapes and progressively elongate the embryo

To examine adherens junction dynamics and correlate them with muscle twitching every few seconds, we used a lattice lightsheet microscope (Planchon et al., 2011). We imaged the *C. elegans* adherens junction marker DLG-1::RFP alone (Fig. 1B), or DLG-1::GFP together with a histone HIS-24::mCherry fusion expressed in muscles (Fig. 1C).

It revealed that embryos first bent laterally for a few seconds at the 1.6-fold stage (Fig. S1B). These movements became progressively more sustained and regular beyond the 1.8-fold stage, when embryos appeared to rotate along their anterior-posterior axis within the eggshell. Typically, the lateral epidermal cells alternated between positions next to the eggshell and at the embryo fold (cyan and yellow dotted lines, respectively, in Figs. 1B, S1C). Labeling muscle nuclei and junctions revealed that muscle contractions, evidenced by nuclear displacement, caused compression of adjacent lateral epidermal cells (cells marked by asterisks in Figs. 1B). Moreover, junction squeezing on one side coincided with stretching on the contralateral side (Fig. S1C).

The cause for rotations became evident when tracking the distance between nuclei in distinct muscle rows (Figs. 1A’). Analysis revealed anti-correlated movement between muscle nuclei in diagonally opposite rows (dorsal left vs. ventral right), explaining the sequential squeezing of left and right lateral cells (Fig. 1C-E; Fig. S1C-D). Cross-correlation analysis tracking lateral epidermal cell deformations delineated partially overlapping anatomical regions in the embryo that deformed independently: the head (H1-H2), the body (H2-V6) (Fig. S2A, B, D), suggesting that different muscles contract independently. Focusing on the 1.8/2-fold stage time window, when recording was easier, we found that embryos spent on average 34% of the time rotating inwardly and 36% outwardly (Fig. S2E-E”; N= 8), demonstrating equal rotation in both directions (for definition of inward/outward rotations, see Fig. 1 legend). The slight 6-degree angle of muscles relative to the anterior-posterior axis (Moerman and Williams, 2006) explains how localized contractions generate a torque inducing embryo rotations.

To assess the impact of rotations on embryonic elongation, we measured the size of adjacent lateral epidermal cells (yellow line in Fig. 2A). We found that their combined AP-length varied between valleys during inward rotations and peaks during outward rotations (Fig. 2B; Fig. S2G-G’), with a period of 41“6 seconds between peaks (Fig. 2C). Remarkably, the total lateral epidermis AP-length progressively increased over time with an elongation rate of approximately 3.1%”1.4 per rotation (Fig. 2D). Meanwhile, the sum of all DV-oriented lateral junctions remained constant over the same period (Fig. 2B). At the cellular level, the head lateral cell H1 underwent less deformation than the body lateral cells (V1-V5) (Fig. 2E-F). This analysis further revealed that muscle and epidermal deformations were coordinated (Fig. S2H-H’).

**Figure 2.**
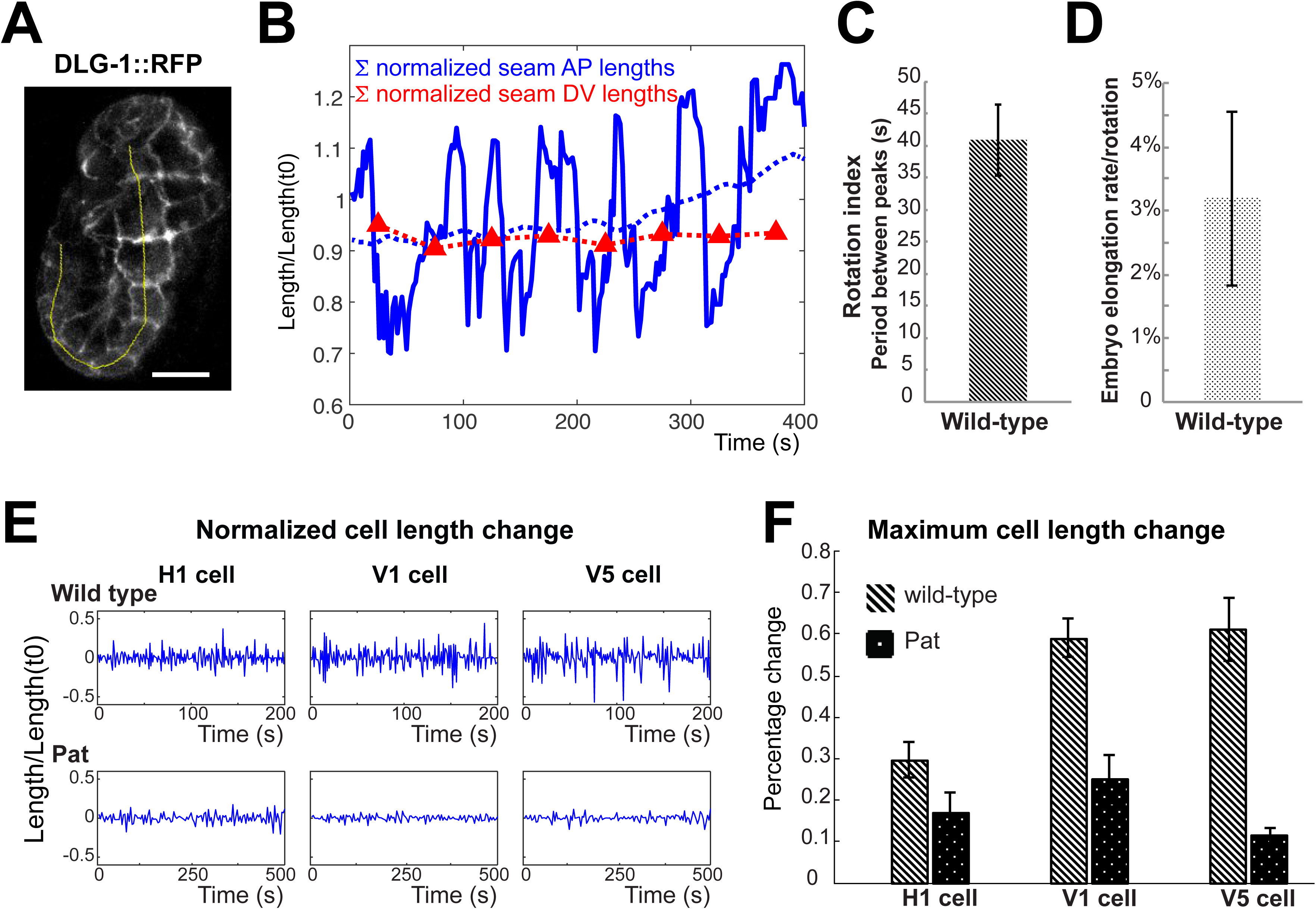
The repeated mechanical inputs stimulate embryo elongation. **(A)** Lightsheet still image of a 1.8-fold embryo expressing DLG-1::RFP (strain ML1652; see Movie S6). Scale bar 10 µm. **(B)** Normalized total length of the left lateral epidermis (thick blue) over time in (a). The blue dotted line shows the corresponding running length average over 100 seconds using the *movmean(time)* from MatLab; the red dotted line shows the normalized sum of all DV junction for the same embryo averaged over 50 second intervals. **(C)** Average time elapsed between major peaks such as in (B) marking the AP-oriented extension of lateral cells (N=8 embryos). **(D)** Average embryo elongation rate relative to major peaks (N=8 embryos). **(E)** Tracking of the apparent AP-oriented normalized length changes for the lateral cells H1, V1 and V5 in wild-type and muscle defective Pat embryos as determined from lightsheet movies (see Movie S7 for *unc-112(RNAi)*). **(F)** Maximum AP-oriented length changes for H1, V1 and V5 epidermal cells in wild-type and *unc-112(RNAi)* embryos. All embryos with their anterior up. See also Fig. S2. Here and in subsequent figures, Pat embryos correspond to *unc-112(RNAi)* embryos.

To examine whether muscle contractions induce shape changes, we abrogated their activity by knocking-down *unc-112*, a typical Pat gene (Rogalski et al., 2000). In 1.8/2-fold embryos, *unc-112(RNAi)* embryos minimally changed lateral cell shapes (Fig. 2E-F), leading to a blurred correlation in shape changes among adjacent lateral cells (Fig. S2C-D). Moreover, these embryos rotated, on average, only 1% of the time (n = 5; Fig. S2F). Hence, muscle contractions cause both transient epidermal shape changes and embryo rotations.

In conclusion, beyond the 1.8-fold stage, the length of lateral epidermal cell oscillates due to muscle contractions, gradually increasing in a polarized manner.

### Muscle contractions exert tension on AP-oriented junctions

The results above indicate that lateral epidermal cells exhibited more folding during so-called inward rotations (Fig. 3A) and more stretching during outward rotations (Fig. 3B). To systematically assess these traits, we quantified the ratio between junction outlines and their smoothened length, or roughness (Fig. 3C). Higher roughness values characterize compressed junctions, whereas lower values indicate stretched junctions. We further correlated roughness with a position during the rotation, taking the V1 lateral cell center as a reference in radial coordinates (Fig. 3D).

**Figure 3.**
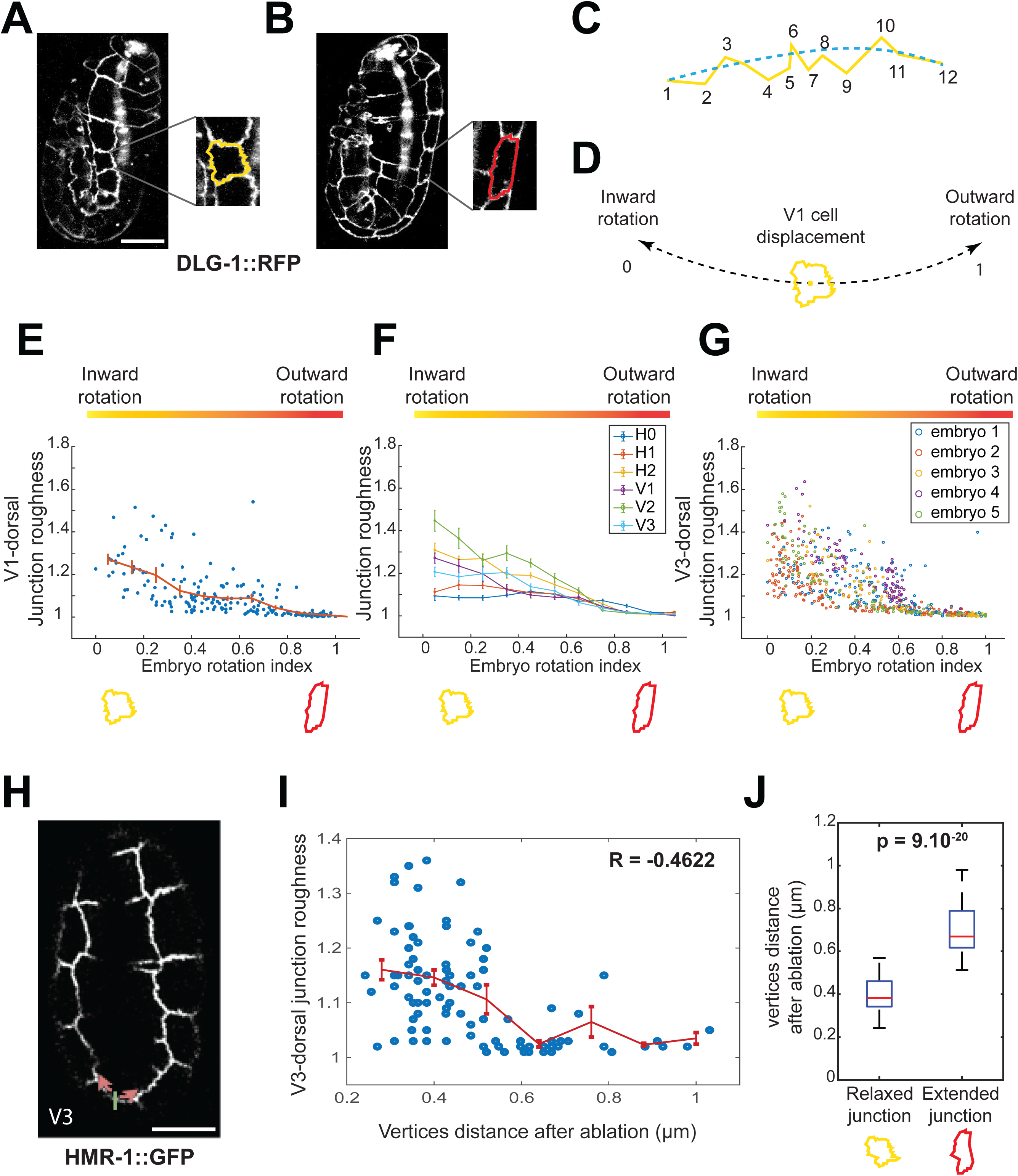
Membrane status during embryo rotations. **(A-B)** Lightsheet micrographs of a wild-type embryo expressing DLG-1::RFP (strain ML1652) showing an inward (a) or outward (b) position, highlighting the folded V1 lateral cell junction (a; yellow line) or the extended V1 lateral cell junction (b; red line). **(C)** Yellow line, actual junction; blue dash line, smoothened junction. **(D)** Embryo rotation index: the black dash curve, represents the V1 cell (yellow) displacement range: position 0, most extreme position during an inward rotation; position 1, furthest position during an outward rotation. **(E-G)** Junction roughness as a function of the embryo rotation index shown for the junctions between the V1 lateral cell and its dorsal epidermal neighbor (E), the lateral cells H0 – V3 and their dorsal epidermal neighbors with a cell-by-cell display (F) or with a display by embryo for the V3 lateral (G); Fig. S3A shows the equivalent roughness index for DV-oriented junctions. Yellow and red cells below the x-axis are as in (A-B). **(H)** 1.8-fold wild-type embryo expressing HMR-1::GFP (strain LP172) used for laser ablation experiments. Short green line, laser ablation area; pink arrows, directions of junction recoil after laser ablation. **(I)** Junction recoil size after laser ablation as a function of junction roughness. R, correlation coefficient. **(J)** Junction recoil size in the first frame after laser ablation (see Fig. S3BC for kymographs). Yellow and red cells are as in (E-G). All embryos oriented with the anterior up. See also Fig. S3.

This procedure indeed confirmed that roughness for AP-oriented junctions increased during inward rotations, and decreased during outward rotations. This trend was observed across all examined lateral epidermal cells in five embryos, with head lateral cells exhibiting a shallower junction roughness slope (Fig. 3E-G). Importantly, DV-oriented junctions remained mostly smooth regardless of their position during rotations (Fig. S3A).

As AP-oriented junctions were more stretched in outward positions, we sought to test whether their tension increased. Using a multi-photon microscope we ablated AP-oriented junctions between the V3 lateral epidermis and the dorsal epidermis in 1.8/2-fold embryos, when rotations are sustained (Fig. 3H). Out of all trackable ablations, AP-oriented junctions recoiled on average by 0.7 µm in 20 embryos rotating outwardly, but only 0.4 µm on average for 66 inwardly rotating embryos, and furthermore junction roughness prior to ablation confirmed that junctions with lower roughness tended to recoil more (Fig. 3I-J; Fig. S3B-C).

We conclude that muscle activity repeatedly induces tensional pulses on AP-oriented junctions.

### E-cadherin concentration at adherens junctions changes over time

So far, we have described how muscle activity influences junction changes at the embryo scale, and specifically promotes lengthening of AP-oriented junctions, thereby reinforcing a pre-existing planar polarity set by the actomyosin cytoskeleton (Vuong-Brender et al., 2017). To explore the kinetics of junctional molecules at a molecular scale, we focused on the core adherens junction component, E-cadherin/HMR-1.

We first assessed E-cadherin concentration by measuring the mean fluorescence intensity of a HMR-1::GFP knockin (Marston et al., 2016) over junction length, keeping consistent imaging settings. In wild-type embryos, HMR-1::GFP fluorescence increased until the 2-fold stage (Fig. 4A-C), then tended to plateau, with only a slight decrease beyond the 3-fold stage. In the head, the E-cadherin signal increased more rapidly in AP-oriented than in DV-oriented junctions, reaching a significantly higher plateau (Fig. 4B). In the body, no apparent difference was observed between AP- and DV-oriented junctions (Fig. 4C). Strikingly, we observed the same pattern in *unc-112(RNAi)* knockdown embryos, both before and after the usual onset of muscle activity (1.8-fold) (Fig. 4A-C).

**Figure 4:**
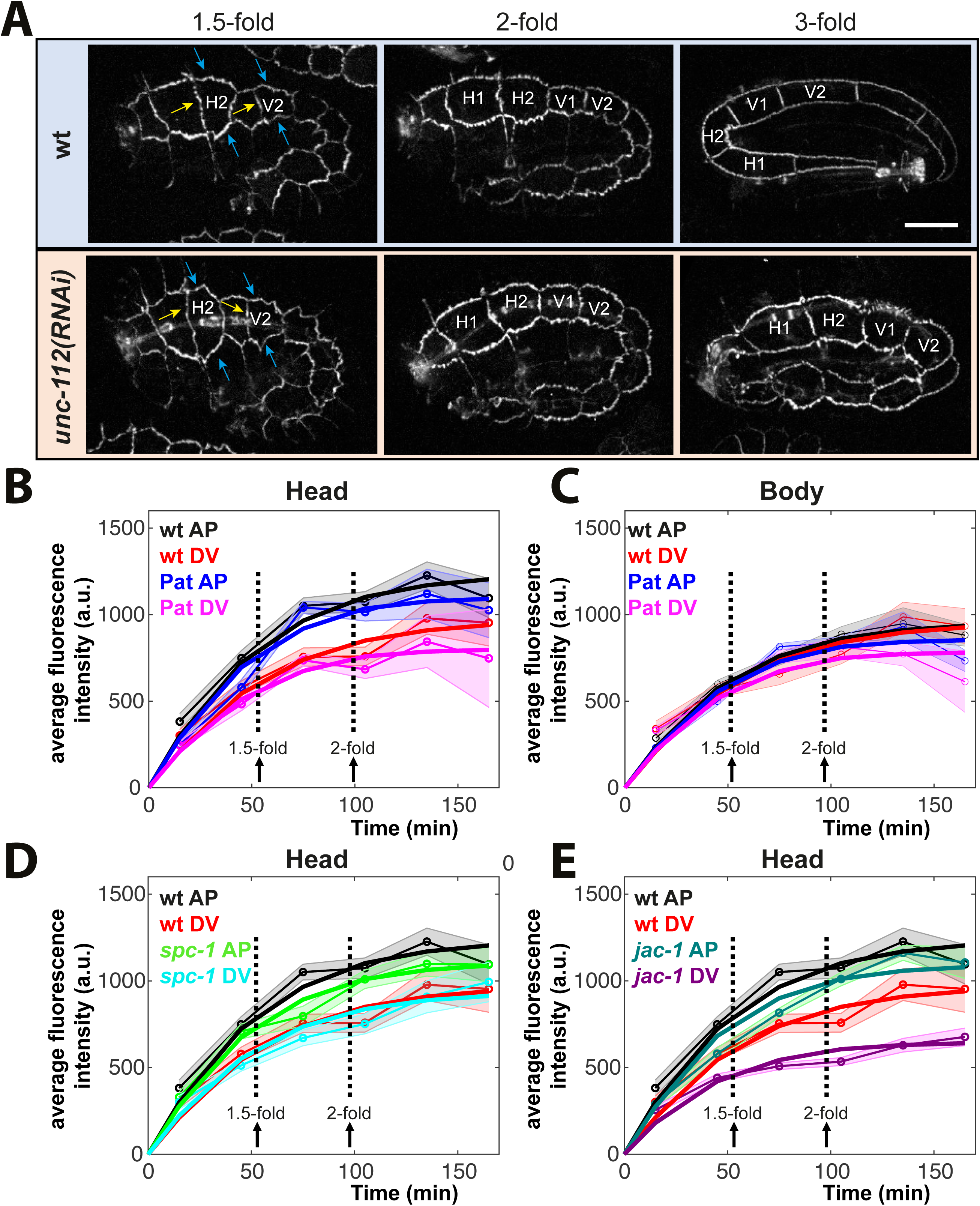
E-cadherin/HMR-1 levels increase until the 2-fold stage along junctions. **(A)** Lateral surface spinning-disk micrographs showing the fluorescence of the CRISPR E-cadherin/HMR-1::GFP (strain LP172) marker at three different stages; settings are identical for the three panels. Blue and yellow arrows indicate the anterior-posterior (AP) and dorso-ventral (DV) junctions, respectively, that were quantified in panels b-c with a focus on the cells H2 and V2; scale bar, 10 µm. Pictures were taken from paralyzed and staged embryos; the stages refer to control embryos and the equivalent time in Pat/*unc-112(RNAi)* embryos. **(B-E)** Quantification of HMR-1::GFP fluorescence using pictures such as those in (A) for the H2-dorso/ventral epidermis (denoted AP) or H1-H2 interface (denoted DV) (B, D-E) in the head, the V2-dorso/ventral epidermis (denoted AP) or V1-V2 interface (denoted DV) (C) in the body, for wild-type (B-E; same values used in B, D-E), Pat (B), *spc-1(RNAi)* (D) and *jac-1(RNAi)* (E) embryos. Shadowed areas, standard errors of the mean. All embryos with their anterior to the left.

In conclusion, the distinction of E-cadherin concentration between AP- and DV-oriented junctions was apparent only in the head, and overall, the addition of new E-cadherin molecules did not directly result from muscle activity. However, since AP-oriented junctions lengthen, it implies that more E-cadherin molecules must be incorporated along AP-oriented compared to DV-oriented junctions.

### Muscle activity promotes E-cadherin turnover during elongation

To refine our analysis, we investigated E-cadherin turnover, first by tracking E-cadherin exocytosis. Using Total Internal Reflection Fluorescence (TIRF) microscopy, we observed potential E-cadherin exocytotic events (Fig. S4). However, acquiring quantitative data was impractical due to TIRF imaging constraints. Consequently, we shifted our focus to Fluorescence Recovery After Photobleaching (FRAP).

To investigate HMR-1/E-cadherin turnover by FRAP, we photobleached an entire junction between two vertices before muscle activity (1.4/1.5-fold stage) and once muscles become active (1.8/2-fold stage), then monitored fluorescence recovery (Fig. 5A-B). Bleaching the entire junction aims to eliminate diffusion effects along the junction. In wild-type embryos, we observed similar signal recovery kinetics at both AP- and DV-oriented junctions, with higher recovery at the 1.5-fold than at the 2-fold stage (Fig. 5C; Fig. S5A, C). The higher recovery in early embryos likely results from the continuous E-cadherin increase at junctions (Fig. 4), but does not explain the higher increase along head AP-oriented junctions. In *unc-112(RNAi)* embryos, E-cadherin fluorescence recovery was equally effective at the 1.5-fold stage but significantly lower at the 2-fold stage compared to control embryos (Fig. 5C; Fig. S5A, C), suggesting more E-cadherin molecules remain at junctions (immobile fraction) in *unc-112(RNAi)* embryos.

**Figure 5:**
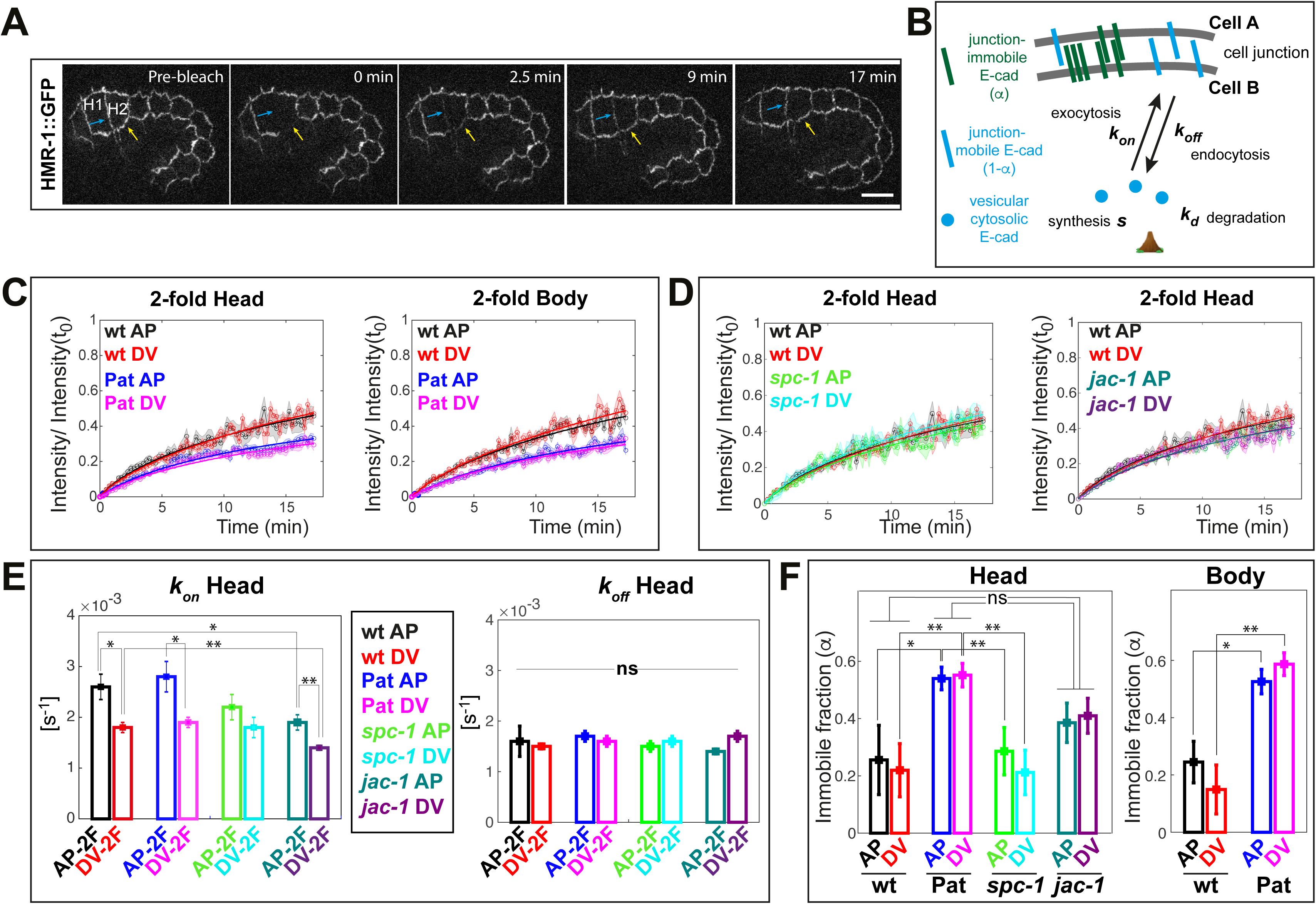
Muscle activity fluidizes junctions to facilitate E-cadherin/HMR-1 turnover. **(A)** Lateral surface spinning-disk micrographs of an embryo at the 1.5-fold stage showing the fluorescence during a FRAP experiment of HMR-1::GFP (strain LP172) for the junctions between the cell H2 and the ventral epidermis (yellow arrows) or with the cell H1 (blue arrows); scale bar, 10 µm. First image, fluorescence before photobleaching; 2^nd^ image, just after photobleaching; next three images, different stages during the recovery process. Note that the embryo elongates during the recovery (see Movie S8). **(B)** Drawing schematizing the exchanges occurring during the FRAP, and the variables *s*, *k_on_*, *k_off_*, *k_d_* and α accounting for HMR-1 dynamics; immobile E-cad molecules a priori correspond to clusters of cis- or trans-interactions. **(C)** HMR-1::GFP recovery profiles at the 1.8/2-fold stage (after muscle activity onset), for the head H2 cell, the body V2 cell in control and *unc-112(RNAi)* (symbolized by Pat) embryos. **(D)** HMR-1::GFP recovery profiles at the 1.8/2-fold stage in control *spc-1(RNAi)* and *jac-1 (RNAi)* embryos (controls are the same as in C). In (C-D) shadowed areas, standard error of the mean. **(E-F)** *k_on_* and *k_off_* (E), and immobile fractions (F) estimated from the fit shown in (C-D), the data in Fig. 4 and the system of equations (1-3, see text; genotype color code between *k_on_*/*k_off_* sub-panels). All embryos oriented with the anterior to the left. See also Fig. S5 for values in the body and at the 1.5-fold stage.

To quantitatively describe E-cadherin behavior, we formulated a kinetic model for E-cadherin concentration *C^AP^* and *C^DV^* on AP- and DV-oriented junctions. Our goal was to thoroughly depict the nature of E-cadherin exchanges along junctions. This model integrates the cytoplasmic E-cadherin reservoir *R* and the main processes regulating E-cadherin turnover, namely: the exocytosis (*k^AP^_on_* and *k^DV^_off_*) and endocytosis (*k^AP^_off_*) rates at AP- and DV-oriented junctions, the synthesis (*s*) and degradation ((*k_d_*) rates of cytoplasmic E-cadherin (Fig. 5B). To account for the higher immobile fraction (α^*AP*^ and α^*DV*^) observed when muscles are non-functional (Fig. 5C), we wrote the E-cadherin kinetics starting at the 2-fold stage as follows (see Methods for the general case):

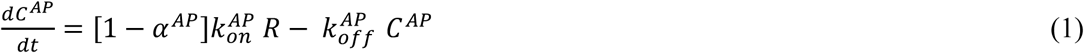

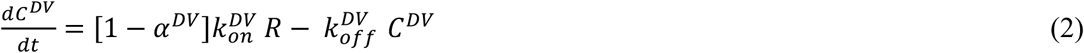

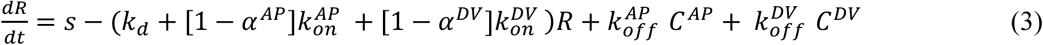

By fitting the data of E-cadherin concentration (Fig. 4BC) and FRAP curves (Fig. 5C), we estimated the values of the endo/exocytosis rates and of the immobile fractions (α^*AP*^ and α^*DV*^), fixing *s* and *k_d_* (see Methods for details). Our data suggest that *k_on_* and *k_off_* values were in the same range, irrespective of the genotype, stage and junction orientation (Fig. 5E; Fig. S5B, D). Furthermore, *k_on_* appeared significantly higher for AP-oriented compared to DV-oriented junctions of the head beyond the 1.8-fold stage (Fig. 5E), accounting for the higher plateau observed for E-cadherin concentration at that stage (Fig. 4B). In the body, *k_on_* for AP-oriented junction was lower compared to head ones (Fig. S5D). Strikingly, at the 2-fold stage, the immobile fraction in the head was higher by a two-fold factor in muscle-defective compared to control embryos, where it exceeded 50% (Fig. 5F). In summary, E-cadherin exocytosis rate appeared higher along head AP-oriented junctions, and its turnover was reduced in muscle-defective embryos everywhere. Of note, the *k_on_* and *k_off_* values reported here correspond to averages over the cell and may hide local variations when the embryo rotates.

The reduced E-cadherin turnover observed in *unc-112(RNAi)* embryos could reflect either their paralysis or lack of elongation. To distinguish between both hypotheses, we examined the kinetics of E-cadherin in embryos defective for SPC-1/α-spectrin, which arrest at the 2-fold stage (Norman and Moerman, 2002), yet can twitch (Lardennois et al., 2019). FRAP analysis of *spc-1(RNAi)* embryos revealed a profile quite similar to that of wild-type embryos in the head (Fig. 5D-F, S5A-B), indicating that muscle paralysis, rather than the lack of elongation, affects E-cadherin turnover in *unc-112(RNAi)* embryos.

Up to this point, we have formulated the kinetic model in terms of concentrations without explicitly considering junction length. To incorporate it into the kinetic equations, we expressed the concentration for AP-oriented junction *C^AP^ = N^AP^/L^AP^*, where *L^AP^* is the AP-oriented junction length, and *N^AP^* is the total number of molecules on the junction. By substituting *C^AP^* and *C^DV^* in equations (1-2), we can write the following equations:

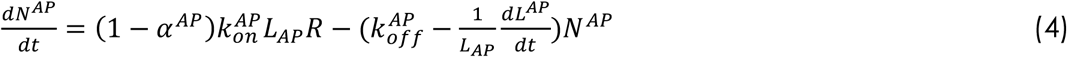

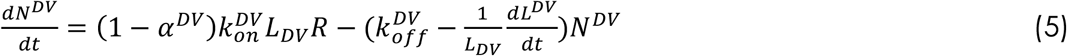

This expression predicts an increase in E-cadherin exocytosis rates as AP-oriented junctions lengthen, with a corresponding decrease in endocytosis since 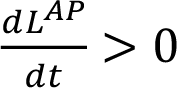. Conversely, endocytosis will increase as DV-oriented junctions shorten since 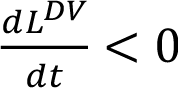. In summary, our analysis shows that muscle activity enhances E-cadherin mobility, facilitating junction fluidity. Additionally, the study shows that E-cadherin concentration remains approximately stable once muscles start contracting (Fig. S5E), implying that junction lengthening and exocytosis promotion go in parallel, while shortening has the opposite effect. This dynamic system maintains concentration homeostasis during morphogenesis.

### A mechano-chemical model accounting for junction elongation

The work presented so far focused on the consequences of muscle activity at the embryo scale, then on the description of E-cadherin kinetics. Since muscle activity and E-cadherin ultimately results in mechanical forces, we sought to develop a mechano-chemical model linking macroscopic tissue deformations to the average E-cadherin molecular kinetics. To this aim, we modelled one lateral epidermal cell as a parallelepiped of sides *l_z_*, *l_y_* and *l_z_*, where *l_x_* and *l_y_* respectively correspond to cell lengths along the AP and DV axis (Fig. 6A). We assumed that the parallelepiped mechanically behaves as an incompressible 3D active viscous fluid (Mayer et al., 2010; Prost et al., 2015) whose deformations take place along the three principal directions only. Additionally, we assumed no friction forces with the surrounding environment as tissues do not move relative to each other at that stage (Sulston et al., 1983).

**Figure 6:**
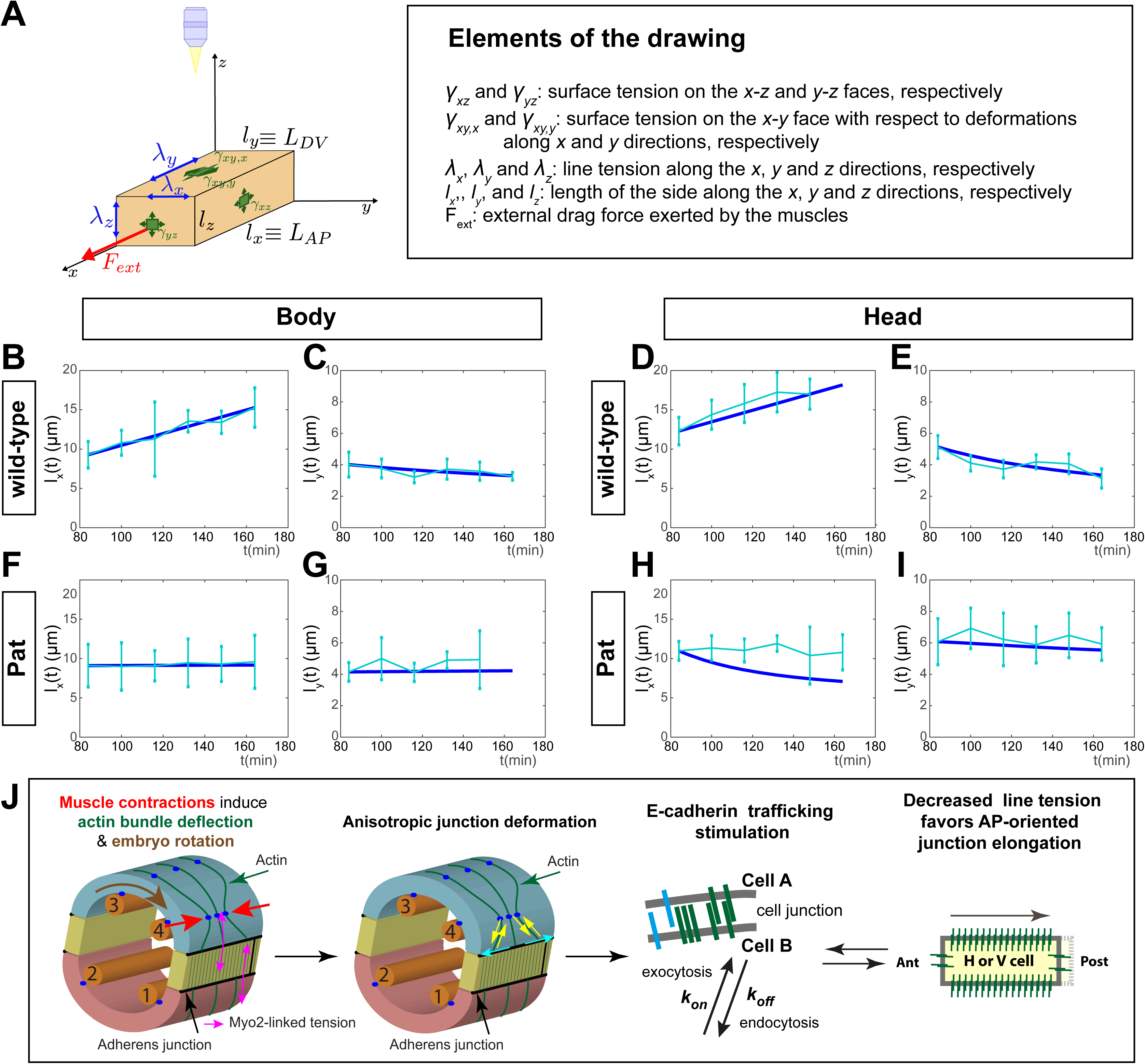
A mechano-chemical model predicting muscle-dependent junction elongation. **(A)** Schematic representation of the model (left drawing) showing the imaging plane, with a description of the parameters in the box on the right. **(B, C)** Result of the simulations from the mechano-chemical model for the wild-type body cell deformations along *l*_0_(B) and *l*_1_ (C). Mechanical parameters are reported in Table S1, whereas the parameters for E-cadherin kinetics are taken from Fig. 5, with *k^AP^_on_*=*k^DV^_on_* =0.1 min^-1^ and *k^AP^_off_*=*k^DV^_off_* =0.084 min^-1^. Thick blue lines in these panels and the following, simulations; thinner cyan lines, experimental data. **(D, E)** Result of the simulations from the mechano-chemical model for the wild-type head cell deformations along *l*_0_ (D) and *l*_1_(E). Mechanical parameters are reported in Table S1, whereas the parameters for E-cadherin kinetics are taken as in Fig. 5, with *k^AP^_on_*=0.16 min^-1^, *k^AP^_off_*=0.1 min^-1^, *k^DV^_on_*=0.1 min^-1^ and *k^DV^_off_*=0.081 min^-1^. **(F, G)** Result of the simulations from the mechano-chemical model for the *unc-112(RNAi)* body cell deformations along *l*_0_ (F) and *l*_1_ (G). Mechanical parameters are reported in Table S1, whereas the parameters for E-cadherin kinetics are taken as in Fig. 5, with *k^AP^_on_*=0.093 min^-1^, *k^AP^_off_*=0.078 min^-1^, *k^DV^_on_*=0.081 min^-1^ and *k^DV^_off_*=0.084 min^-1^. **(H-I)** Result of the simulations from the mechano-chemical model for the *unc-112(RNAi)* head cell deformations for *l*_0_x (H) and *l*_1_ (I). Mechanical parameters are reported in Table S1, whereas the parameters for E-cadherin kinetics are taken as in Fig. 5, with *k^AP^_on_*=0.15 min^-1^, *k^AP^_off_*=0.09 min^-1^, *k^DV^_on_*=0.11 min^-1^ and *k^DV^_off_* =0.09 min^-1^. See also Fig. S6. **(J)** Model for the influence of muscle contractions on E-cadherin turnover. 1^st^ diagram, muscle contractions induce embryo rotations (brown circular arrow) and deflect actin bundles (red arrows). 2^nd^diagram, as a consequence the stress on junctions increases either perpendicularly (yellow arrows) or along them (cyan arrows). 3^rd^diagram, E-cadherin turnover gets stimulated. 4^th^diagram, line tension favours junction elongation and in turn stimulates the recruitment of additional E-cadherin molecules (see bidirectional arrows between 3^rd^ and 4^th^ diagrams).

The different terms in the stress equation can be divided into passive and active terms (Prost et al., 2015). The passive terms are respectively given by: i) a viscous term characterised by a viscosity coefficient η and ii) a pressure term Δ*P* = *P* − *P*_0_ quantifying the difference between the internal and the external pressure, *P* and *P*_0_, and imposed by volume conservation. The active terms, generated by cytoskeleton remodelling and cadherin adhesion, are respectively given by the sum of the contributions due to: i) the surface tension, i.e., the force per unit of length to deform the surface lying on the *i − j* plane along the *k*-th direction, γ_*ij, k*_, and ii) the line tension, i.e., the force to deform the junctions lying along the λ-th direction, λ_*i*_ (see supplementary theory for details). To take into account anisotropic tension along the DV axis resulting from higher actomyosin activity in that direction (Vuong-Brender et al., 2017), we assumed an increased surface tension in the *x-y* plane.

As discussed elsewhere(Arslan et al., 2024; Maitre et al., 2012; Montel et al., 2021), cadherin concentration promotes contact expansion by locally reducing tensions. Since epidermal E-cadherin localizes only to adherens junctions, we assumed its relative concentration linearly decreases line tension. On the other hand, surface tensions are assumed to be homogeneous, isotropic, and equal for all surfaces except the x-y plane. To take into account anisotropic tension along the DV axis resulting from higher actomyosin activity in that direction (Vuong-Brender et al., 2017), we assumed that surface tension on the *x* − *y* plane with respect to deformation in the DV-direction is increased compared to that characterizing deformation in the AP-direction. Therefore, we imposed *γ_xy,x_* = γ_0_ and γ_*xy,y*_ = μ γ_0_.

The constitutive equations for the stress components along the three principal directions are therefore given by (see Supplementary Theory for the detailed derivation):

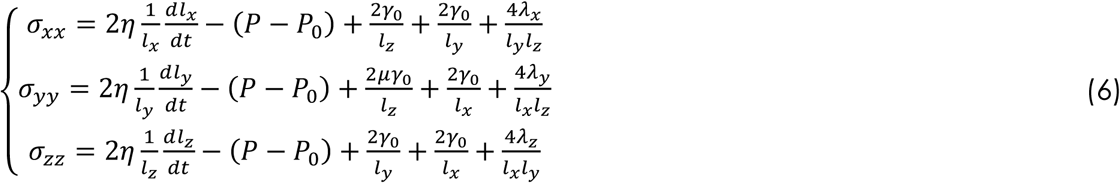

By assuming a constant average drag-force *f* exerted by muscles on the cell, the force balance reads:

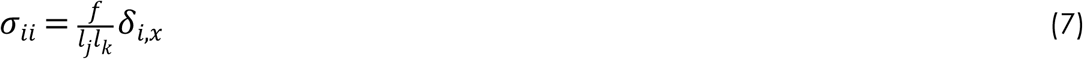

where *i, j, k* = *x, y, or z* and δ_*i,x*_ is equal to 1 if *i* = *x* and 0 otherwise.

We first used this model to understand to what extent the difference in E-cadherin endo/exocytosis and the anisotropy in surface tension induced by actomyosin along the DV direction (Vuong-Brender et al., 2017) contribute to deformations. As shown in Fig. S6A-C, a difference of E-cadherin exocytosis along the AP or DV direction generally favors cell thinning along the z-direction. Yet, a higher E-cadherin concentration along one direction reduces line tension and thus favors lengthening along that direction and shortening along the z-direction (Fig. S6A-C). The effect of actomyosin cables instead leads to contraction along the DV direction, thus elongation along the z-direction (Fig. S6D).

We next sought to quantitatively assess the contributions of different forces to wild-type and muscle-defective embryos. To determine the drag-force induced by muscle activity, we derived the average wild-type AP speed of elongation from experimental observations (Fig. 6B and 6D), then fitted the remaining line and surface tension parameters, namely λ_0_, γ_0_ and μ, together with the cadherin conversion factor ε. For simplicity, we assumed the mechanical parameters in the head to be uniformly higher than in the body by a factor κ to reflect its greater stiffness (Vuong-Brender et al., 2017). As shown in Figs. 6B-E, S6E-F, the mechano-chemical model accurately replicated DV-oriented junction shortening and AP-oriented junction lengthening in the head and body, predicting also shortening along the z-axis consistent with embryo diameter reduction. To assess the predictive power of the model in the absence of external drag-forces, we simulated the force-balance equations in Pat mutants, using for simplicity the basal mechanical parameters fitted in the wild-type case and the cadherin kinetics previously obtained (Figs. 4, 5). Notably, the model successfully predicted the scarcity of deformations in the absence of muscle-induced drag-forces in both body and head cells (Figs. 6F-I, S6G-H).

In conclusion, the combined action of the muscle drag-force and non-muscle myosin defines a polarization in the mechanical deformations. Moreover, by leveraging FRAP experimental outcomes on E-cadherin kinetics (Fig. 5), we could robustly estimate the values of mechanical parameters such as the ratio between surface tension and viscosity (Mayer et al., 2010), the magnitude of the anisotropy generated by the actomyosin along the DV direction, the difference in stiffness between the body and the head (Vuong-Brender et al., 2017) and, importantly, the line tension and its link with E-cadherin concentration (Fig. S6I-J) without any direct mechanical measurement, thus avoiding fatal disruptions of embryos.

### E-cadherin trafficking levels modulate elongation

The mechano-chemical model predicts that E-cadherin trafficking levels may affect elongation. To explore this, we first knocked-down p120-catenin, a regulator of E-cadherin endocytosis in other systems (Bulgakova and Brown, 2016; Ishiyama et al., 2010; Iyer et al., 2019; Sato et al., 2011). In p120-catenin/JAC-1 knockdown embryos, E-cadherin concentration and *k_on_* along head AP and DV junctions were significantly reduced (Fig. 4E, Fig. 5E). Moreover, the immobile fraction was higher than control levels, although not as high as Pat levels (Fig. 5F). As *k_off_* rates are similar across genotypes (Fig. 5E), they probably reflect only the mobile fraction; hence, the larger immobile fraction in *jac-1(RNAi) embryos* likely reflects reduced endocytosis. Consistent with these mild effects and previous findings showing that JAC-1 seems dispensable for morphogenesis (Pettitt et al., 2003), we observed that *jac-1(RNAi)* young L1 larvae hatched at 230 µm, slightly but significantly shorter than the 240 µm wild-type L1 hatchlings (Fig. S6K).

Second, we used our mechano-chemical model to predict how reduced endocytosis or exocytosis affects elongation. The simulations indicate that reducing endocytosis even by a 10-fold factor, but not exocytosis, moderately decreases elongation without completely blocking it (Fig. S6L-O). Notably, the rather mild effects observed in *jac-1(RNAi)* embryos align with these predictions and with estimates of the muscle drag force-to-E-cadherin line tension ratio, which is approximately 18 in the head and 2.5 in the body (Fig. S6P).

In summary, our findings suggest that elongation is mainly driven by the muscle drag force, with a positive contribution from E-cadherin trafficking.

## DISCUSSION

Through light-sheet microscopy, laser ablation, FRAP analysis, and modelling, we have explained how an external force within the embryo stimulates the polarized extension of AP-oriented adherens junctions and the addition of more E-cadherin along those junctions.

Unlike in other systems (David et al., 2010; Fernandez-Gonzalez et al., 2009; Maitre et al., 2012; Martin et al., 2009; Rauzi et al., 2008), we argue that the muscle-driven junction elongation operating in *C. elegans* does not involve junctional myosin II since it is largely absent from junctions (Molnar et al., 2024). Furthermore, there are no apical actomyosin flows (Vuong-Brender et al., 2017), as in the aforementioned studies. It also does not rely on a mechanically-induced change in tissue planar polarity or the mobilization of internal membrane reservoirs, as observed elsewhere (Aigouy et al., 2010; Lecuit and Wieschaus, 2000). Indeed, laser ablation experiments showed tension mainly oriented along the DV axis (Vuong-Brender et al., 2017). Additionally, the possibility of membrane reservoirs within the epidermis seems unlikely due to the limited time available to build reservoirs between epidermis arising and muscle contractions (Sulston et al., 1983). Moreover, membrane reservoirs would not account for the specific extension of AP-oriented junctions.

Instead, we propose that muscle contractions have two effects (Fig. 6J). First, FRAP analysis of E-cadherin turnover demonstrated that muscle contractions enhance junction fluidity. Our findings concur with published data on the timescales of junction fluidization (20-40 seconds) (Bambardekar et al., 2015; Campas et al., 2024; Clement et al., 2017; Mongera et al., 2023). It may for instance lower the binding energy of E-cadherin cis- and trans-dimers to reduce the E-cadherin immobile fraction (Campas et al., 2024). Second, a mechano-chemical model recapitulating the observed *in vivo* deformations predicted that an E-cadherin imbalance in the AP direction reinforces deformation in that direction though line tension. Combined with our mathematical analysis of FRAP data, it explains why more E-cadherin molecules are incorporated along AP-oriented junctions as they lengthen. Furthermore, mechano-chemical predictions and a direct genetic test indicate that disrupting E-cadherin trafficking can impact junction dynamics and embryonic elongation.

At the time scale of each muscle contraction (seconds), two factors could contribute to the preferential E-cadherin recruitment along AP-oriented junctions. First, lightsheet microscopy suggests that during muscle-induced embryo rotations, AP-oriented junctions alternate between a flabby state and a more tensed state. Past observations indicate that membrane tension stimulates exocytosis and reduces endocytosis (Sheetz and Dai, 1996). Second, AP-oriented junctions are submitted to orthogonal tension resulting from the DV-oriented anisotropic tension (Vuong-Brender et al., 2017), which should be reinforced when muscles deflect dorso-ventral actin filaments (Lardennois et al., 2019). This tension is known to be disrupted in Pat mutants since actin filaments of lateral cells reorient in the AP direction in Pat mutants (Gillard et al., 2019). Tensile force perpendicular to junctions generally increases E-cadherin levels by stabilizing trans-dimers (Engl et al., 2014; Kale et al., 2018; le Duc et al., 2010; Yonemura et al., 2010). We suggest that both factors exert their effect locally, either when junctions are on the outer edge and/or when muscles contract (Fig. 6J). We do not exclude additional input from actin-binding proteins; for instance, the WAVE complex promotes E-cadherin turnover in the *C. elegans* intestine (Sasidharan et al., 2018).

Although as argued above, the process described herein differs from other systems relying on repeated inputs, such as *Drosophila* gastrulation, germband extension or dorsal closure (Bertet et al., 2004; Fernandez-Gonzalez et al., 2009; Martin et al., 2009; Rauzi et al., 2010; Solon et al., 2009), some elements are common, which should represent general features in mechanobiology. First, the repeated nature of the mechanical input modifies the tension exerted on junctions, and progressively induces junction shrinking or lengthening. In *Drosophila*, the mechanical inputs correspond to regular actomyosin flows with a period of ≈90 sec, with additional influence from neighboring tissues (Collinet et al., 2015; Lye et al., 2015). In *C. elegans*, the tensile inputs correspond to muscle contractions with a period of ≈40 sec which are transmitted to the epidermis via the muscle-epidermis attachments (this work). Second, those mechanical inputs are controlling E-cadherin trafficking [(Levayer et al., 2011), this work]. Third, the anisotropy is in part originating from global anisotropy in the embryo, reinforced by the mechanical process. In the *Drosophila* germband, the global anisotropy is controlled by segmentation genes that feed on transmembrane proteins coupled to heterotrimeric G-protein, and ultimately on actomyosin activity (Kerridge et al., 2016; Lavalou et al., 2021; Pare et al., 2019; Pare et al., 2014). In *C. elegans*, earlier tension and stiffness anisotropy in the embryo (Vuong-Brender et al., 2017), combined with muscle activity (this work), orient the polarity of junction lengthening.

The present results reveal how muscles drive gradual junction lengthening, and, along with previous findings (Lardennois et al., 2019; Zhang et al., 2011), provide a comprehensive view of *C. elegans* elongation. Furthermore, together with other studies, they further illuminate how multiple tissue types mechanically coordinate their morphogenesis (Aigouy et al., 2010; Collinet et al., 2015; Lardennois et al., 2019; Lye et al., 2015; Norman and Moerman, 2002; Zhang et al., 2011). Much like past genetic work has laid the groundwork for reprogramming any cell type into specific tissues, our findings can facilitate the mechanical bioengineering of functional organ-on-chip systems.

## Supporting information

Includes the supplemental theory and 6 suppelemntal figures

## ACKNOWLEDGEMENTS

The authors thank Ananyo Maitra, Grégoire Michaux, François Robin, Raphaël Voituriez and Alpha Yap for critical comments on the manuscript, all past members of the Labouesse, in particular Sophie Quintin for an illuminating discussion, members of the Robin lab for suggestions, as well as Sophie Le Cann, Guillaume Salbreux and Raphaël Voituriez for fruitful discussions. We thank the IBPS Imaging Facility for advice. This work was supported by grants from the Agence Nationale pour la Recherche (grants #ANR-18-CE13-0008-01 and ANR-22-CE13-0037), the European Research Council (grant #294744) to ML. ML warmly thanks the MPI-PKS Visitor’s Program for supporting his stay in Dresden.

## AUTHORS’ CONTRIBUTIONS

ML and XY performed the lightsheet imaging using a setup designed by NM, LR and GM; XY did all the lightsheet image analysis with input from JP and TF. She also performed the laser ablation. TF performed the FRAP analysis and data modelling. SG and SRM elaborated the mechano-chemical modelling. SWG gave advice while hosting ML. ML conceived the project and wrote the first draft, which was finalized by ML, XY, TF and SG, with input from all authors. XY and ML assembled figures, TF wrote the supplementary FRAP modelling material, SG wrote the supplementary theory section.

## DECLARATION ABOUT THE USE OF AI

We used AI technology to improve the readability and concision of the text, carefully checking for the meaning of each proposal for scientific relevance.

## STAR Methods

### Animal strains, conditions of maintenance

*C. elegans* strains used in this study are listed in Table S2. Animals were maintained at 20°C, unless stated otherwise, on Nematode Growth Medium (NGM) agar plates as described in (Brenner, 1974). N2 Bristol was used as the wild type strain.

### Engineering knockin markers

Knockin DLG-1::GFP and HMR-1::pHluorin::HMR-1 insertions were obtained by the method described in (Dickinson et al., 2013). For DLG-1, the GFP coding sequence was positioned just upstream of the *dlg-1* stop codon. For HMR-1, the modified worm-specific super-ecliptic pHluorin was amplified from plasmid pSJN788 (Dittman and Kaplan, 2006) (a gift from Dr. Josh Kaplan) and inserted at position S838 within the extracellular domain. A repair template for Designing Homologous Repair was constructed in the pJET1.2 vector (Thermo Fisher Scientific) by the Megawhop method (Miyazaki and Takenouchi, 2002). The repair template included the new sequence to be inserted and 1500 or 500 base pair of homologous sequence on either side of the insertion site. The Protospacer Adjacent Motif (PAM) was mutated (protein sequence was not changed) in the repair template to prevent the plasmid and the repaired genomic locus from being cleaved by Cas9. The small guide RNA (sgRNA) sequence specific to *dlg-1* or *hmr-1* was inserted in the pDD162 vector (Dickinson et al., 2013) that also encodes Cas9 under a germline promoter. The primer and SG sequences used in this study are described below.

*dlg-1* sgRNA sequence: CCAATTTCATCTAATGACG

*dlg-1* Megawhop primers:

Forward: ACACATGGCATGGATGAACTATACAAACGTCATTAGATGAAATTGGATA

Reverse: CCCCAAAAAGCAAAAGCAGGAAAATTAAAATGCAAGTATTTGTAAGGTG

*hmr-1* sgRNA sequence: AAAAGGGTTGTAGGCTGTG

*hmr-1* Megawhop primers:

Forward: TGTGGGAAGCCACGTGTGACAGTAACTCAAGTAAAGGAGAAGAACTTTTCACTGG Reverse: ATGCAATGATTCAACGAGTCGACTTTGTATAGTTCATCCATGCCATGTG

The repair template and Cas9/sgRNA plasmids were co-injected into both gonads of wild-type adult hermaphrodites with the roller marker pRF4 and the red pharynx marker pCFJ90 [Pmyo-2::mCherry] (Frokjaer-Jensen et al., 2008). Rollers or animals with a red pharynx from the F1 progeny were picked and checked by PCR to find positive hits after egg-laying. One successful knock-in was found in 40 F1 animals for DLG-1::GFP, and one for HMR-1::pHLuorin::HMR-1. The absence of a mutation at the site of insertion was verified by sequencing. Homozygous *mc103[dlg-1::gfp]* embryos and larvae were healthy and behaved like wild-type controls. On the other hand, homozygous *mc118[hmr-1::pHLuorin::hmr-1]* animals showed 60% lethality, yet could be maintained in a homozygous state, and embryos that passed the 2-fold stage could develop successfully to give progeny.

### RNAi feeding

We used the Ahringer feeding RNAi library to inactivate *unc-112*, *spc-1* or *jac-1* (Fire et al., 1998; Kamath and Ahringer, 2003). Wild-type L4 larvae were fed for 48 hours before transferring them to a fresh RNAi plate. 24 hours later, the embryos between 1.5/2-fold were selected for imaging. In parallel, the terminal phenotype of embryos was scored to ensure RNAi efficiency and found to generate 95% two-fold arrested embryos. For *jac-1(RNAi)*, mothers were grown on a fresh RNAi plate for 48 hours. The efficiency of *jac-1(RNAi)* was tested by exposing *hmp-1(fe4)* animals to RNAi; we found that the hatching rate after 48 hours was 90% for *hmp-1(fe4)* grown on control bacteria versus 5% on *jac-1(RNAi)* producing bacteria, which matches previously reported results (Pettitt et al., 2003).

### Lightsheet imaging

Lightsheet imaging was done using a microscope built up by N. Maghelli and L. Royer in MPI-CBG, Dresden (Royer et al., 2015) based on the lattice lightsheet microscope developed by Eric Betzig and collaborators (Chen et al., 2014). The objective used for imaging was 40x, and embryos were maintained at 25°C. We first selected embryos undergoing the process of dorsal epidermal cell fusion (1.5-fold stage) and started a pre-acquisition time-lapse with 2-minute time intervals until embryos actively twitched or rotated. Continuous imaging was then started. During pre-acquisition and acquisition, we selected embryos with lateral epidermal cells parallel or nearly parallel to the camera. Embryos were scanned from one side to the other side, with a focal plane depth of 0.5 µm. We used DLG-1 rather than HMR-1/E-cadherin as an adherens junction marker, because in pilot tests the signal appeared less prone to bleaching under continuous imaging. For one-colored movies of the strain ML1652 (see Table S2 for strain genotypes), the exposure time was from 3-10 ms and the acquisition process was continuous imaging. Due to the frequency of muscle contractions, we acquired a complete stack in approximately 0.4/0.45 seconds and limited movies to 500 seconds to avoid phototoxicity. For two-colored movies of embryos from the strain ML2616, the acquisition was with 2 second time intervals. For all time-lapses, control overnight movies with a 2-minute time interval were run to ensure that the imaging protocol was not phototoxic for embryonic development, and discarded acquisitions in which embryos stopped development, which was very rarely the case.

### Lightsheet movie analysis

To calculate the changes of cell aspects, the vertices of lateral epidermal cells from SPIM movies were tracked in Fiji. For control embryos the tracking was finished manually and the original positions of vertices were saved. For *unc-112(RNAi)* embryos, the tracking of POI (Points of Interest) was done by a Fiji plugin, *CE Time lapse analysis of POI*, which was created by Julien Pontabry. Briefly, the initial POI was compared in consecutive frames to find the potential point with exact or similar surrounding background using a cross-correlation function. After gathering the original positions of all POIs, we used *shape inserter* in MATLAB to circle the position of each POI back on the SPIM movies to make sure that the original tracking of the POI was correct. After confirming so, we calculated the apical area and the perimeter of the cell by a pre-existing MATLAB function. function [geom, iner, cpmo] = polygeom (x, y). Function website: https://fr.mathworks.com/matlabcentral/fileexchange/319-polygeom-m

The length of the cell was calculated as the total length from the mid-point of the two most anterior vertices passing the central position of all the vertices to the mid-point of the two most posterior vertices of the same lateral epidermis. The positions of these three points were calculated in MATLAB by using the original positions of the vertices of a lateral epidermis. The length was calculated by:

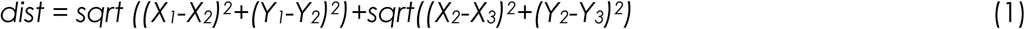

The total length of all lateral epidermis is the sum of the length of all the lateral epidermal cells. To track the distance between two nuclei, the positions of two selected muscle nuclei from each row of muscles was tracked in Fiji at each time point, and the distance was calculated in MATLAB using equation (1). To simplify the analysis of embryo rotation by skeletonizing the AP-length (see Fig. S2E-E’’), we somewhat arbitrarily drew two lines reflecting the time points when the ventral junction of the V1 lateral cell was nearing the inside edge of the embryo during an inward rotation, or when its dorsal junction was nearing the outside edge during an outward rotation.

To measure the roughness of junctions during embryo rotation, Lightsheet movies were loaded in MATLAB frame by frame. In each frame, the position of every vertex was recorded by the MATLAB function *ginput* (Graphical input from mouse or cursor). Subsequently, the position of each vertex was smoothened by the MATLAB function *smooth* with the exponential smooth option. Based on the actual position of the smoothened position of each point, the total length and the smoothened length of each junction were calculated by adding up the distance between each vertex. Based on the tracking of junctions during rotation, a smoothened length of the junction was calculated by MATLAB automatically using the following equation:

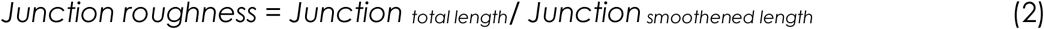

### Laser ablation experiment

Laser ablations of adherens junction were conducted on a Leica SP8 multi-photon microscope using a 63×/N.A. 1.4 oil-immersion objective on the strain LP172. Embryos older than 1.7-fold were selected for ablation at the junction between V3 lateral epidermis and dorsal cells. For each set of ablations, we acquired three frames pre-ablation and 50 frames post-ablation with a 488 nm laser. The ablation was done by the multi-photon laser with a wavelength of 904 nm at an average laser power of 1700 mW using a Region of Interest (ROI) of 5 nm x 0.08 nm, diagonal to junctions. The exposure time for pre- and post-ablation was 0.5 second; it was 0.27 second for ablation. As the laser ablation experiment was conducted on rotating embryos, which can result in a loss of focus of the ablated junction as early as the first frame post-ablation - it was the case for approximately 100 embryos - we always measured the distance between the two vertices of the ablated junction in the first frame post-ablation. Since it took 1.25 seconds for the microscope to change from a 904 nm ablation laser mode to the 448 nm laser imaging mode, the distance between the two vertices represented the junction recoil size 1.25 seconds after laser ablation. The pixel size was 0.08615 µm under the laser ablation setting; accordingly, the outward rotating junctions presented an average recoiling of 80 pixels and the inward rotating junctions an average recoiling of 46 pixels (see Fig. 3I). Among the 116 successfully ablated embryos for which we managed to track the post-ablation junctions, 30 of them actually died after ablation or elongated abnormally, as shown by our systematic long post-ablation recording, and were excluded. The analysis of recoil size was done in FiJi. The average intensity of selected junctions was measured first in the pre-ablated imaging. After laser ablation, pixels with an intensity lower than 10% of the average intensity of the pre-ablation junction were considered as the recoil and the recoil size was measured.

### Single molecule imaging and analysis of fusion events

Single molecule imaging was conducted on a Nikon Total Internal Reflection Fluorescence (TIRF) Microscope. Embryos older than 1.7-fold were selected for imaging. To image the HMR-1::pHLuorin::HMR-1 construct (strain ML2773), a 488 nm laser was used. The initial laser power was set up at 100 mW. The exposure time was 80 ms with continuous imaging. Before imaging, embryos were bleached for three rounds until the histogram showed that the peak of imaging intensity decreased to between 7,500 and 10,000. Each round lasted for 30 seconds, with 80% laser power. The incident laser was set up at 0° to bleach through the whole embryo. Between each round of bleaching, the embryo rested for 45 seconds without any imaging to avoid high photo-toxicity. After bleaching, the rotations of the embryos were imaged with the incident light set up at the optimal degree, usually between 64° and 67.5°.

After imaging, one or more sub-portions of a movie presenting an AP- or DV-oriented junction in sharp focus and in a stretched position during rotation was selected for analysis. The duration of these short movies was usually 4-10 seconds (40-130 frames 80 ms/frame). By using a MATLAB interface, a region surrounding the selected junction was masked. From each time series, the difference in intensity between two frames *D(t)* was calculated to extract potential fusion events. *D(t)* was defined as follows:

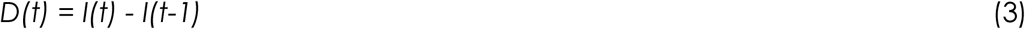

where *I(t)* is the frame at time *t* and *I(t-1)* is the frame at the previous time. *D(t)* was filtered by a gaussian filter with sigma=1 to average out noise. Then a threshold was set up for all the sub-portions of a movie, based on a representative frame. The threshold was the same for all sub-movies belonging to the same movie and was estimated with respect to the background of a region outside the junction as following:

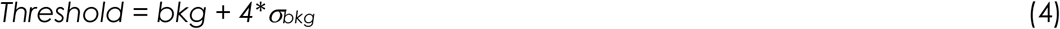

where *bkg* was the average value of the background and *α_bkg_* was its standard deviation. After thresholding, fusion events were automatically registered and analyzed. A fusion event was identified as a super-threshold spot with a size bigger than 4 pixels. For each fusion event the integral fluorescence intensity and the position of the spots were recorded. Moreover, the distance between the two vertices of the selected junction in each frame was measured.

### E-cadherin junctional signal imaging and quantification

E-cadherin/HMR-1::GFP fluorescence signal was measured using the CRISPR allele *hmr-1(cp21)* (strain LP172) (Marston et al., 2016). We imaged embryos with a spinning-disk Zeiss microscope Axio Observer.Z1 using a 100X oil immersion objective, a 491 nm laser at 15% power with an exposure time of 150 ms controlled by the Metamorph software, as described elsewhere (Lardennois et al., 2019). We acquired z-stacks of 50 focal planes spaced 0.4 µm apart. The same settings were used for all genotypes. From the original maximum z projection created by Fiji, we subtracted the background estimated by image averaging on a rolling ball radius of 50 pixels. The E-cadherin signal along the junctions was measured using the Fiji line profile tool with a width of 3 pixels and recording the average intensity.

To get clean images of fluorescence profiles (Fig. 4) we immobilized embryos by oxygen deprivation obtained by mounting eggs in a medium rich in bacteria. We staged wild-type embryos based on their body length. We staged *unc-112(RNAi)* based on their body length until the 2-fold stage, since their elongation rate was normal until that stage (Lardennois et al., 2019). Beyond the 2-fold stage, *unc-112(RNAi)* showed progressive deformation of the lateral junctions in time lapse movies (data not shown), and we staged them based on those deformations. In our hands, *spc-1(RNAi)* by feeding was not as strong as by injection and resulted in embryos that had an almost normal elongation rate until the 1.8-fold stage, yet did not extend much beyond the 2/2.2-fold length. We had previously showed that their DV-oriented junctions keep shrinking once they have reached the 1.8-fold stage (see Supp Mat of (Lardennois et al., 2019)) and we used that feature to stage *spc-1(RNAi)* embryos.

Note that since the average HMR-1::GFP intensity slightly increases while the AP-oriented junctions lengthen and the DV-oriented junctions shrink beyond the 2-fold stage (Fig. 4), the total number of HMR-1 molecules along AP-oriented junctions must increase while it must decrease or remain constant along DV-oriented junctions (Fig. S5E).

### FRAP imaging and quantification

FRAP experiments were performed on the CRISPR allele *hmr-1(cp21)* (Marston et al., 2016) (strain LP172) using the iLaS^2^ module integrated to the same spinning disk system described above. For each embryo, single or multiple full junctions were photo-bleached by using 100% power of the 491 nm laser using the Fly mode. We acquired 10 z-stacks spaced 0.4 µm apart and centered on the focal plane of interest prior photobleaching, then captured 12 z-stacks every 15 s for up to 1040 s after bleaching, with an exposure time of 80 ms and a laser power of 15%. We photobleached full junctions to exclude molecule diffusion during fluorescence recovery in subsequent image analysis.

Every 4-d image file was treated by maximal z-projection for all the time points using Fiji. The background was then subtracted for all time points, using the dedicated function implemented in Fiji with a rolling ball radius of 50 pixels. After image pre-treatment, the mean fluorescence intensity along each photobleached junctions was recorded by using a manual segmented line with 7 pixels width in Fiji. The recovery curves used to estimate the kinetic parameters were obtained using the equation:

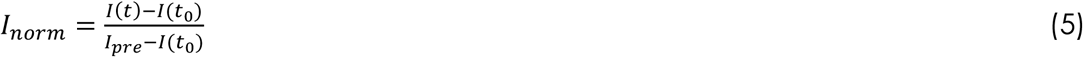

where *I(t)* is the fluorescence intensity at the time point *t*, *t_0_* being the first time point after photobleaching and *I_pre_* the average intensity computed on the last three time points preceding the photobleaching. Correction for post-FRAP photobleaching has been unnecessary.

### Fitting E-cadherin behavior

The curves describing E-cadherin concentration during elongation (Fig. 4) and FRAP recovery curves (Fig. 5C) are ruled by the same kinetics. Before the 2-fold stage the E-cadherin signal is out of equilibrium since the fluorescence at junctions monotonously increases. Indeed, the FRAP curves acquired at the 1-5 stage are affected by the global increase of the signal, such that, to properly fit the data, we used a time dependent model and not a steady state solution. According to a simple first order kinetic, the concentration of molecules is controlled by the following basic processes: the synthesis of E-cadherin molecules in a cell with a rate *s*, its degradation rate in the cytoplasm *k_d_*, the rate of E-cadherin exocytosis *k_on_* and the rate of E-cadherin endocytosis *k_off_.* If we call *C^AP^* and *C^DV^* the concentration of E-cadherin on one AP- or DV-oriented junction, we can write the following equations:

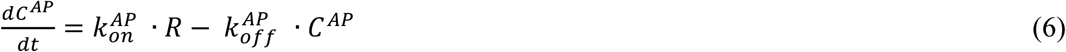

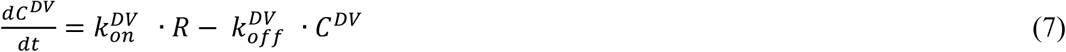

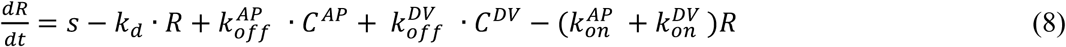

where *R* is the concentration of the cytosolic reservoir. This model can fit the wild-type data at different stages of elongation including the FRAP curves for the two stages (1.5-fold and 2-fold) with a constant value of the *on* and *off* rates. On the other hand, the wild-type and *unc-112(RNAi*) FRAP recovery curves at 2-fold have a similar half-time, but different values for the asymptote, and cannot be fitted together with the global behavior of the E-cadherin during elongation. For that reason, we decided to include a new parameter, the immobile fraction that we called α to account for a portion of molecules contributing to the total concentration but not to the FRAP recovery curves. Since the difference is manifest at the 2-fold stage, we chose to introduce the immobile fraction at that stage. Hence, we modified the equations (5-7) as follows:

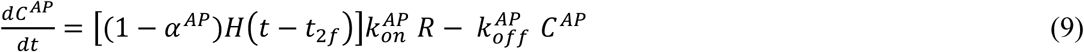

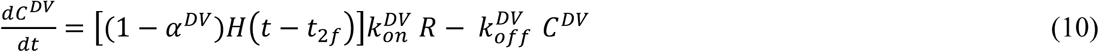

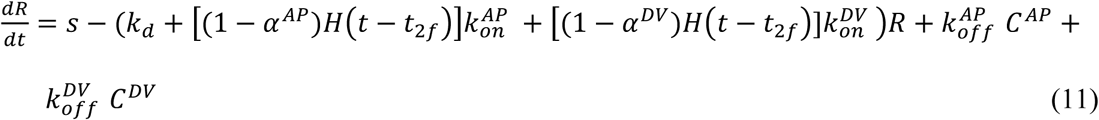

where *H*(*t* − *t*_2*f*_) is the Heaviside step function that is equal to 1 for *t* ≥ *t*_2*f*_ and zero otherwise, where *t*_2*f*_ states the time at which the 2-fold stage is reached. The advantage of this formulation is the presence of an immobile fraction in explicit form in the time-dependent model.

To reduce the number of free parameters, we fixed the values of *s* and *k*_(_. The value of *k*_(_, combined with the endocytosis rate *k_off_*, affects the transient duration of the fluorescence behaviors before the plateau. From the FRAP curves we can have an estimation of *k_off_* as the inverse of the half time of the curves ^(Sprague et al., 2004)^. Among the different cells and genotypes we can see that the inverse of the curve’s half-time is of the order *k_off_* ∼1*x*10^−3^ *s^−1^*. We then used this value to pre-simulate the system of Eqs. (9-11), explore the order of magnitude of the degradation rate and account for the equilibration time of the E-cadherin signal. It resulted in *k_d_* being of the same order of magnitude as *k_off_* with an optimum value of *k_d_* = 2.5*x*10^−3^ *s*^−1^, which we used.

The synthesis rate, *s*, is a scale factor accounting for the observed E-cadherin concentration at steady state (*C_ss_*), which, according to Eqs. (9-10), can be written as 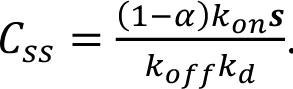. The observed fluorescence level of the plateau is of the order of ∼10^3^ (Fig. 4B-C). However, this value is arbitrary, since we do not have a calibration between the fluorescence level and the number of molecules. We chose to attribute the same value of ***s***=2 *s*^−1^ for cells in the head, body and among the different genotypes. This choice implies that the parameter *k_on_* has to be in the ∼10^-3^ *s^−1^* range. To estimate the 6 parameters 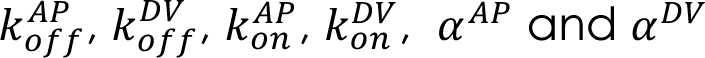, we fitted the average FRAP recovery curves along the AP and DV junctions and the average E-cadherin intensity profiles along the AP and DV. Every cell considered for FRAP has four junctions and we frapped two of them in Fig. 5A. Indeed, two pools of molecules contribute to the recovery curves: the cytoplasmic reservoir and the non-bleached molecules recycled from the non-frapped junctions. To define whether or not the pool of E-cadherin on junctions was negligible compared to the cytoplasmic one, we performed a FRAP experiment in which all junctions from two neighboring cells were bleached (Fig. S5F). We observed a roughly twice lower recovery curve compared to experiment such as in Fig. 5 in which only two junctions are bleached (Fig. S5G). Therefore, we solved simultaneously Eqs. (9-10) twice, once for bleached junctions with initial fluorescence concentration being zero and once for the non-bleached junctions with initial fluorescence concentration being equal to the pre-bleached concentration. In summary, for each bleached cell in an embryo, we solved a kinetic equation for each of the four junctions and a kinetic equation Eqs. (11) for the cytosolic reservoir.

Assuming that Eqs. (9-11) have constant kinetic parameters, we first fitted them at the 1.5-fold stage, estimating the four parameters 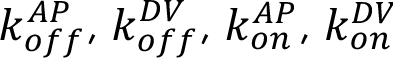. Since at the 2-fold stage we introduced α^*AP*^ and α^*DV*^, to reduce the degree of freedom, we imposed an upper limit to *k_off_* as 1.7*x*10^−3^ *s^−1^*, which represents the highest value of the *off* rates observed at 1.5-fold (Fig. 5). We estimated the error on the parameters by fitting the data with their standard error and taking the half of variability interval has the error. The best fit parameters were obtained by minimalizing the following cost function:

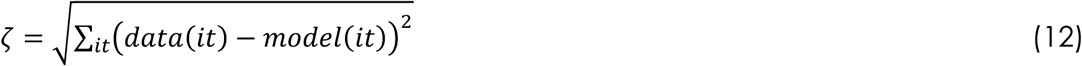

where *data(it)* is the value of the data at the time point *it* and *model(t)* is the solution of the model at the same data point *it*. To minimize the cost function, a Covariance Matrix Adaptation Evolution Strategy (CMA-ES) algorithm was used. The algorithm has an available MATLAB code at https://www.mathworks.com/matlabcentral/fileexchange/52898-cma-es-in-matlab. Eqs(9-11) have been solved numerically with a custom made code written in MATLAB.

All MATLAB scripts used for the present analysis are available upon request.

### Statistical tests

All pairwise comparisons were done using unpaired Student’s test. The number of samples for each experiment is reported in Table S3. The correlation coefficient for Fig. 3I was calculated using the function *corrcoef()* in MathLab.

## Supplementary Figure legends

**Supplementary Figure 1: The alternating pattern of muscle contractions sequentially modify the apparent AP length of the lateral epidermis**

**(A-A)** Major cellular features of an embryo. Left side of a 1.8-fold embryo showing the positions of the underlying muscles (orange lines in A). The cross-section shows the organization of the actin filaments or actin bundles in the epidermis (A’); internal organs were omitted for clarity. Enlarged scheme of the mechanically coupled epidermis-muscle tissues (A’’) highlighting the hemidesmosome-like structures connecting the epidermis to the extracellular matrix apically and ventrally, and the focal adhesion complexes anchoring myofilaments to the muscle plasma membrane. **(B)** Still pictures from a lightsheet pre-acquisition movie of a 1.6-fold ML1652 embryo shortly after the fusion of the dorsal epidermal cells showing the first major deformation of the lateral epidermis at the level of the V1 cell (nomenclature of lateral cells in the left-most picture; see Movie S3). Timing in the upper left corner. **(C)** Still pictures from a lightsheet movie of a 1.7-fold ML1652 embryo showing its dorsal side with the left (blue arrows) and right (yellow arrows) V1-V3 lateral cells being alternatively compressed or relaxed/slightly extended, presumably coinciding with the alternating contraction pattern of the underlying left and right muscles (see Movie S4). Timing in the upper left corner. **(D)** Still pictures from a two-color lightsheet movie of strain ML2616 showing that lateral cell compression coincides with a muscle contraction (see Movie S5). The distance (in µm) along the AP axis between two muscle nuclei (yellow arrows; Muscle Nuclei Distance, M.N.D.), and the AP length of the lateral cells V1 and V2 (white arrows; see Movie S5) are indicated below. Timing in upper right corner. Scale bars, 10 µm. All embryos oriented with the anterior up.

**Supplementary Figure 2: Elongation-defective mutants affect lateral cell shape changes and embryo rotations**

**(A)** Two by two comparisons of the apparent AP-oriented deformations underwent by the lateral cells H1 in the head, V1 and V2 in the body and V5 in the posterior body, as observed in a lightsheet movie of a wild-type embryo expressing DLG-1::RFP (strain ML1652). Note that the head and body cells were generally not deformed simultaneously. **(B-D)** Systematic cross-correlation of the apparent AP-oriented deformations underwent by the H0-V6 lateral epidermal as deduced from lightsheet movies of wild-type (B) and *unc-112(RNAi)* (C) embryos (strain ML1652). The pattern of lateral cell deformation defines two semi-independent cell groups in a wild-type embryo with the tail cell V6 potentially constituting a third group (D). Blue circle, head group; red circle, body group; green circle, potential tail group. These groups were lost in *unc-112(RNAi)* embryos. **(E-E”)** Normalized length of the lateral epidermis and rotation pattern for one particular wild-type embryo (strain ML1652). Blue line apparent normalized length of the lateral cells on one side (E), with correction for absolute length (E’) and skeletonization to symbolize the rotation status from inspection of movies (E’’; see Methods). Value #1” represents outward rotations as values above the upper yellow threshold line in (E’); value #-1” represents inward rotations as values below the bottom yellow threshold line in (E’); value #0” represents relaxed positions. **(F)** Skeletonized rotation status of an *unc-112(RNAi)* embryo (strain ML1652). **(G-G’)** Half-length of a DLG-1::RFP embryo (G), corresponding to yellow lines in the images shown in (G’); the numbers along the curve (G) correspond to specific panels in (G’). Note the perfect match between inward rotations (images 2 and 5) and the shortened lateral epidermis length, or between outward rotations (image 4) and its increase. Time points (G) are separated by 0.4 second. Scales bars, 10 µm. All embryos oriented with the anterior up. **(H-H’)** Still picture from Movie S2 highlighting two specific muscles (circled) and three specific lateral epidermal cells (H2, V1, V2) (H), and distance over time between those muscles (cyan line) in parallel to the AP length of H2-V1-V2 (yellow line) in (H). Note that both tracing are in synchrony, illustrating how muscle and the epidermal movements are coordinated.

**Supplementary Figure 3: Junctions positioned outward experience higher tension**

**(A)** The roughness of DV-oriented junctions was calculated as outlined in Fig. 3C, here for the junction between the head lateral cells H1 and H2 (left panel) and for the body lateral cells V1 and V2. **(B)** DIC picture (B1), image of the V3 lateral-dorsal junction prior (B2) and immediately after (yellow arrow in B3) laser ablation, along with a kymograph of the opening (red arrow in B4). In this 1.9-fold embryo (strain LP172); the junction was closer to the eggshell than to the fold. **(C)** DIC picture (C1), image of the V3 lateral-dorsal junction prior (C2) and immediately after (yellow arrow in C3) laser ablation, along with a kymograph of the opening (red arrow in C4). In this embryo (strain LP172), the junction was closer to the fold than the eggshell. Scale bar, 10 µm (B1-B3 and C1-C3) or 1 µm (B4 and C4). All embryos oriented with the anterior to the left.

**Supplementary Figure 4: Strategy to visualize E-cadherin exocytotic events by TIRF**

**(A)** Scheme of the HMR-1protein, and position at which the super-ecliptic pHluorin was inserted in the extracellular domain using CRISPR technology. CA, Cadherin repeat; EGF, Epidermal Growth Factor-like domain; LamG, Laminin G domain. **(B)** Principle of the ecliptic pHluorin fluorescence: we anticipated that the fluorescence of the pHluorin should be quenched within vesicles (dark green in the left panel) until their fusion with the plasma membrane (light green in right panel). **(C-C’)** Total Internal Reflection Fluorescence (TIRF) micrograph after bleaching most fluorescence within the embryo (strain ML2773), so as to statistically measure the fluorescence of unique molecules. The image shows a junction after image treatment. **(C’)** Six consecutive images of the area boxed in (C) separated by 80 ms showing a brief burst of fluorescence likely to represent an exocytotic event (yellow arrowhead). Due to the frequent rotations of the embryos and the limitation of TIRF imaging, which requires imaging a single focal plane, most junctions could not be tracked for a long time, making it difficult to generate enough data for analysis.

**Supplementary figure 5: FRAP parameters at the 1.5-fold stage and in the body**

**(A)** HMR-1::GFP FRAP recovery profiles at the 1.5-fold stage (before muscle activity onset), for the head H2 cell in control, *unc-112(RNAi)* (symbolized by Pat) embryos, *spc-1(RNAi)* and *jac-1 (RNAi)* embryos (controls are the same in both sub-panels). **(B)** Values for HMR-1::GFP *k_on_* and *k_off_* at the 1.5-fold stage inferred from (A). **(C)** HMR-1::GFP FRAP recovery profiles at the 1.5-fold stage for the body V2 cell in control and *unc-112(RNAi)*/Pat embryos. Shadowed areas in (A, C), standard error of the mean. The number of embryos for each genotype is over 12. **(D)** Values for HMR-1::GFP *k_on_* and *k_off_* in the body for control and *unc-112(RNAi)*/Pat embryos at the 1.5-fold stage, inferred from (C), and 2-fold stage (inferred from Fig. 5C). **(E)** Total HMR-1::GFP fluorescence intensity as a proxy for the total number of HMR-1 molecules over time for wild-type embryos in H1-H2 head cells and V1-V2 body cells. Shadow, standard error of the mean. **(F)** Lateral surface spinning-disk micrographs of a control embryo at the 1.5-fold stage showing the fluorescence recovery after photobleaching all junctions of cells V1 and V2 (yellow arrows); images correspond to the control pre-bleach, immediately after the bleach (t_0_), and at two subsequent time points. Scale bar, 10 µm. All embryos oriented with the anterior to the left. **(G)** HMR-1::GFP FRAP recovery profiles of the DV-oriented junction between V1 and V2, comparing 1.5-fold embryos in which two junctions had been bleached (as in Fig. 5) or seven junctions had been bleached (as in panel F, see cyan dotted rectangle).

**Supplementary Figure 6: Controls for the mechanochemical model**

**(A-C)** Phase diagrams showing the sides of the parallelepiped, *l*_0_, *l*_1_ and *l*_2_, respectively, at t=80 minutes as a function of the ratio between the E-cadherin binding (on) and unbinding (off) rates along the AP and DV directions. The remaining kinetic parameters for E-cadherin are given by *s*=150 min^-1^, *k_d_*=0.15 min^-1^, *α^AP^*=0.26 and *α^D^*^V^=0.15, as obtained from FRAP experiments (see Figs. 5 and S5). Mechanical parameters are instead given by *ψ_0_/ι=*0.0012 µm*min^-1^, *λ_0_/ι=*0.067 µm^2^ * min^-1^, *ε/ι*=0.011 µm*pN^-1^*min^-1^ and *µ*=1. **(D)** *l*_0_, *l*_1_ and *l*_2_ as a function of the scaling factor *µ*. All parameters are taken as in panels (A-C) other than *ε* =0 and the rates of E-cadherin binding/unbinding here taken as in the fit of the wild-type body cells, i.e., *k^AP^_on_*=*k^DV^_on_*=0.1 min^-1^ and *k^AP^_off_*=*k^DV^_off_*=0.084 min^-1^. **(E-H)** Results for the fits from the mechano-chemical model matching Fig. 6B-I for the deformations along *l_z_*; mechanical parameters and E-cadherin kinetics parameters are indicated in the legend for Fig. 6 for each condition. **(I)** Distributions of the logarithm for the sets of parameters optimally reproducing the data *π*^∗^ = (*ψ0/11*, *α_0_/11*, *χ/11*, *μ* and *κ*). **(J)** Coefficients of variation for each of the parameters. **(K)** Size of wild-type and *jac-1(RNAi)* young hatchling larvae in a box-plot representation. N≥45 for each genotype; the p value is shown above the box-plot. **(L-O)** Simulations from the mechano-chemical model for the wild-type head cell deformations along the *l_y_* direction (L, N) or *l_z_* direction (M, O) when *k_off_* (L, M) or *k_on_* (N, O) are made to vary from 10% to 100% of the control value (see Supplementary Theory for details). The reason for not varying E-cadherin turnover along the x direction is that the model has been based on the wild-type parameters for *l_x_(t)*. However, given that the cell is incompressible, if, as predicted by the simulations *l_y_(t)* increases when *k_off_* is reduced, it is likely that *l_x_(t)* will increase less than in the control situation. **(P)** Estimate of the relative ratio of the muscle drag force versus the E-cadherin linked line tension. Values correspond to an average over time (symbol *& >_t_*).

## Supplementary Movie legends

**Movie S1**: Lightsheet movie of the strain ML1652 expressing DLG-1::RFP. The still pictures from Fig. 1B correspond to the time points 434-450. Acquisition, one frame every 0.45 s; playing speed, 25 fps.

**Movie S2**: Montage from a two-color lightsheet movie of the strain ML2616 showing the left dorsal side (on the left) and right ventral side (on the right). The still pictures from Fig. 1C correspond to the time points 166-169. Acquisition, one frame every 2 s; playing speed, 25 fps.

**Movie S3**: Lightsheet movie of the strain ML1652 at the very beginning of muscle twitching (so-called pre-acquisition movie, see Methods); note that junction between dorsal epidermal cells can be seen disappearing at the beginning of the movie due the cell fusion process. The still pictures from Fig. S1B correspond to the time points 8-12. Acquisition, one frame every 2 min; playing speed, 2 fps.

**Movie S4**: Lightsheet movie of the strain ML1652 soon after the beginning of muscle twitching The still pictures from Fig. S1C correspond to the time points 122-134 (out of 250). Acquisition, one frame every 0.45 s; playing speed, 25 fps.

**Movie S5**: Two-color lightsheet movie of the strain ML2616. The still pictures of Fig. S1D correspond to the time points 98-102 (out of 300). Acquisition, one frame every 2 s; playing speed, 25 fps.

**Movie S6:** Lightsheet movie of the strain ML1652 corresponding to the still picture from Fig. 2A. Acquisition, one frame every 0.45 s; playing speed 25 fps.

**Movie S7:** Lightsheet movie of a 2-fold muscle-deficient embryo (ML1652 strain fed with *unc-112(RNAi)* bacteria) corresponding to the still pictures from Fig. S2D and the tracing of Fig S2F. Acquisition, one frame every 0.45 s; playing speed 10 fps.

**Movie S8:** Maximum projection from a FRAP movie of the strain LP172 showing HMR-1::GFP recovery, and corresponding to the still pictures from Fig. 5A. The video starts when the embryo is approximately 1.5-fold, ending when it is almost 1.7-fold and starts to show muscle contractions. Acquisition, one frame every 15 s; playing speed 20 fps.

## Supplementary Tables

**Table S1.**
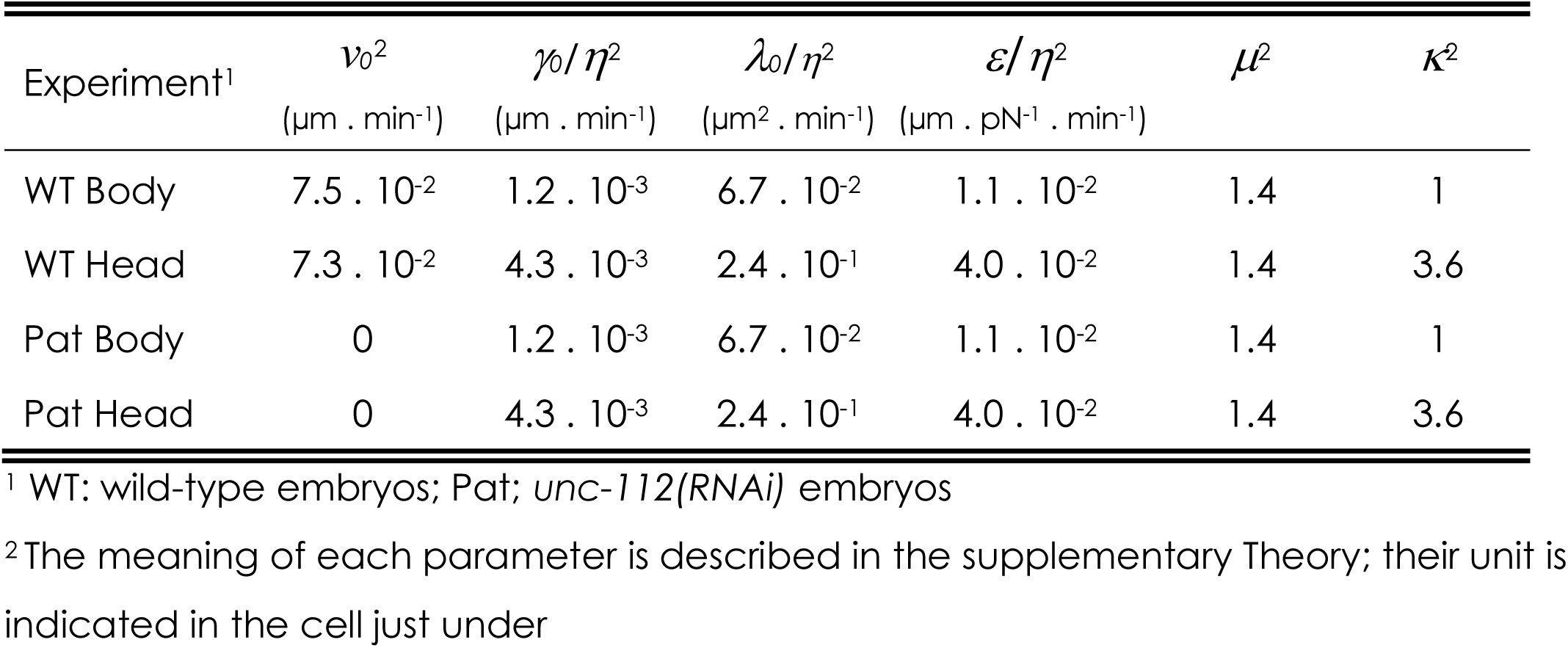
Average parameter values obtained from the search algorithm described in the Supplemental Theory section

| Experiment<sup>1</sup> | $v_0^2$<br>($\mu\text{m} \cdot \text{min}^{-1}$) | $\gamma_0/\eta^2$<br>($\mu\text{m} \cdot \text{min}^{-1}$) | $\lambda_0/\eta^2$<br>($\mu\text{m}^2 \cdot \text{min}^{-1}$) | $\varepsilon/\eta^2$<br>($\mu\text{m} \cdot \text{pN}^{-1} \cdot \text{min}^{-1}$) | $\mu^2$ | $\kappa^2$ |
| --- | --- | --- | --- | --- | --- | --- |
| WT Body | $7.5 \cdot 10^{-2}$ | $1.2 \cdot 10^{-3}$ | $6.7 \cdot 10^{-2}$ | $1.1 \cdot 10^{-2}$ | 1.4 | 1 |
| WT Head | $7.3 \cdot 10^{-2}$ | $4.3 \cdot 10^{-3}$ | $2.4 \cdot 10^{-1}$ | $4.0 \cdot 10^{-2}$ | 1.4 | 3.6 |
| Pat Body | 0 | $1.2 \cdot 10^{-3}$ | $6.7 \cdot 10^{-2}$ | $1.1 \cdot 10^{-2}$ | 1.4 | 1 |
| Pat Head | 0 | $4.3 \cdot 10^{-3}$ | $2.4 \cdot 10^{-1}$ | $4.0 \cdot 10^{-2}$ | 1.4 | 3.6 |

<sup>1</sup> WT: wild-type embryos; Pat; *unc-112(RNAi)* embryos
<sup>2</sup> The meaning of each parameter is described in the supplementary Theory; their unit is indicated in the cell just under

**Table S2.**
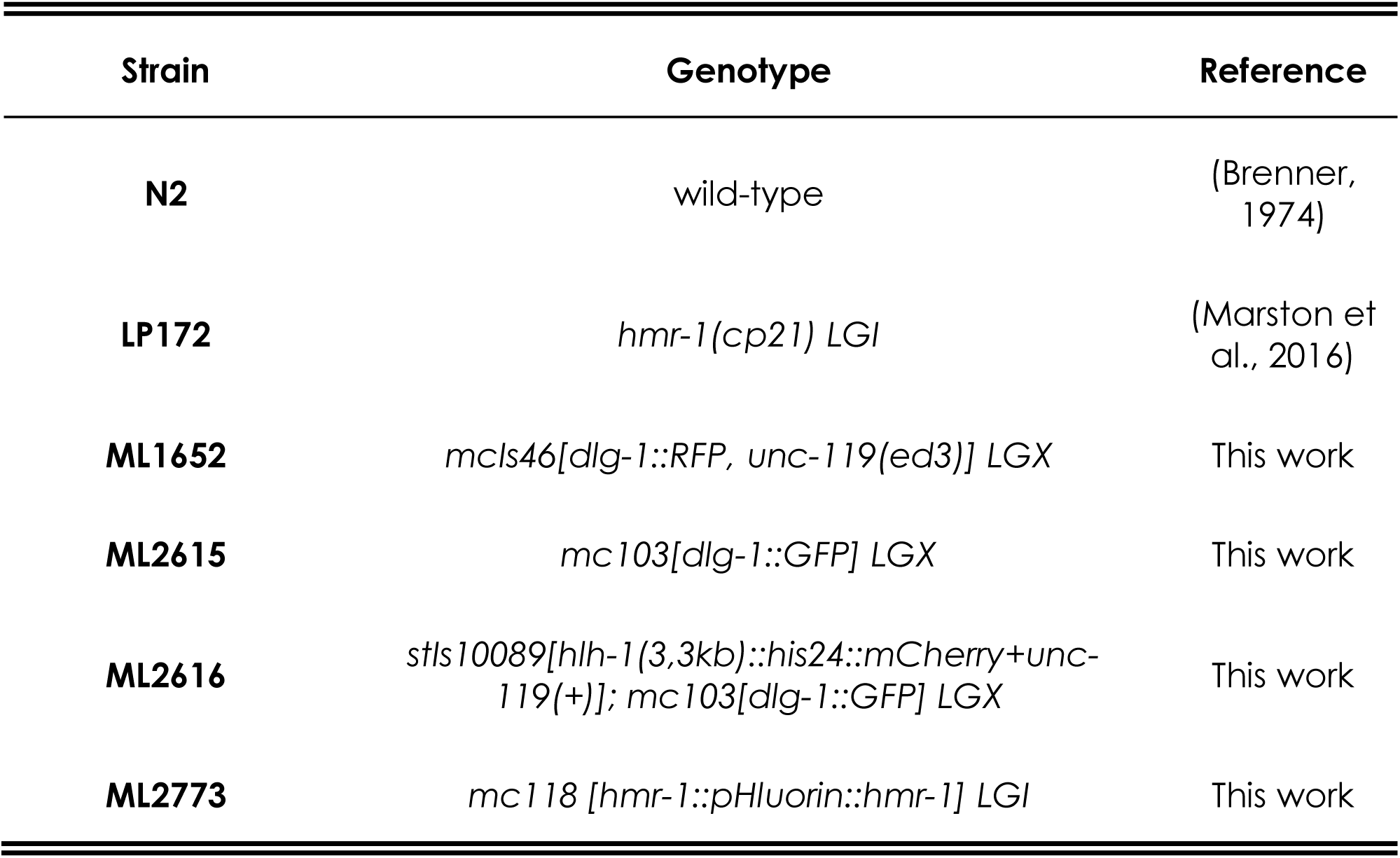
Strains used in this study.

**Table S3.** Sample size

| **Figure** | **Stage<sup>1</sup>** | **wild-type<sup>2</sup>** | ***unc-112(RNAi)*<sup>2</sup>** | ***spc-1(RNAi)*<sup>2</sup>** | ***jac-1(RNAi)*<sup>2</sup>** |
| --- | --- | --- | --- | --- | --- |
| Figure 1E |  | 4 |  |  |  |
| Figure 2C-D |  | 8 |  |  |  |
| Figure 2F |  | 8 | 5 |  |  |
| Figure 3I |  | 86 |  |  |  |
| Figure 4B-E |  | 82 | 61 | 54 | 100 |
| Figure 5E | 1.5 fold head | 20 | 16 |  | 19 |
|  | 1.5-fold body | 13 | 14 |  |  |
|  | 2-fold head | 44 | 12 |  | 45 |
|  | 2-fold Body | 36 | 12 |  |  |
| Figure S5B | 1.5-fold head | 20 | 16 | 15 | 19 |
|  | 2-fold head | 44 | 12 | 25 | 45 |
| Figure S5D | 1.5 fold body | 20 | 16 |  |  |
|  | 2-fold body | 13 | 14 |  |  |
| Figure S5G | 2 junctions | 13 |  |  |  |
|  | 7 junctions | 5 |  |  |  |
| Figure S6K |  | 45 |  |  | 46 |

<sup>1</sup> Refers to the embryonic stage at which the experiment was conducted
<sup>2</sup> Sample size

