## Supplementary material for "Repeated extrinsic and anisotropic mechanical inputs promote polarized adherens junction elongation": Includes the supplemental theory and 6 suppelemntal figures

### SUPPLEMENTARY THEORY

We consider one single cell in the worm embryo beyond the two-fold stage and model it as an active viscous fluid enclosed in a parallelepiped. We define the sides of the parallelepiped as $l_x, l_y$ and $l_z$ respectively and impose the volume, $V_0$, to be conserved. $l_x$ and $l_y$ respectively correspond to the Anterior-Posterior length, $L_{AP}$, and the Dorso-Ventral length, $L_{DV}$, as shown in Fig. 6A. Volume conservation leads to an equation constraining the three sides of the parallelepiped such that:

$$\frac{1}{l_x}\frac{dl_x}{dt} + \frac{1}{l_y}\frac{dl_y}{dt} + \frac{1}{l_z}\frac{dl_z}{dt} = 0. \tag{1}$$

Since experimentally only deformations along the three main axis are observed, we assume that the only non-zero components of the stress tensor are given by the diagonal terms which read:

$$\begin{cases} \sigma_{xx} = 2\eta \frac{1}{l_x}\frac{dl_x}{dt} - f_x \\ \sigma_{yy} = 2\eta \frac{1}{l_y}\frac{dl_y}{dt} - f_y \\ \sigma_{zz} = 2\eta \frac{1}{l_z}\frac{dl_z}{dt} - f_z \end{cases} \tag{2}$$

where $\eta$ is the shear viscosity and $f_i$, where $i = x, y$ or $z$, is the contribution to the internal forces densities to the stress coming from the difference between the internal and external pressure and the line and surface tensions. Since the experimental timescales span over about hundreds of minutes, we assume that elastic contributions to the stress tensor are negligible and therefore passive deformations are characterised only by viscous deformations and internal pressure [1, 2].

We assume that at mechanical equilibrium the force to deform one of the sides is given by the line tension $\lambda_j$ whereas the force per unit of length to deform the surface lying on the $i-j$ plane (where $i, j = x, y$ or $z$ and $i \neq j$) is given by the surface tension $\gamma_{ij}$. Yet, previous works based on laser cuts experiments [2, 3] showed that the system displays anisotropies in surface tensions along the AP and DV direction. To take this into account, we assumed two different surface tensions for the surface lying on the $x-y$ plane according to the direction of deformation, i.e., $\gamma_{xy,x}$ (resp. $\gamma_{xy,y}$) for the surface tension of the surface lying on the $x-y$ plane with respect to deformations in the AP (resp. DV) direction. All other surface tensions, i.e., $\gamma_{xz}$ and $\gamma_{yz}$, are assumed to be isotropic and homogenous. The form of the line tensions instead will be specified in the following.

To compute the contributions coming from the internal pressure and tensions, we define the variation of the mechanical work in the fluid, $\delta W$, as follows:

(3)
$$\delta W = 2\gamma_{xy,x} l_y \delta l_x + 2\gamma_{xy,y} l_x \delta l_y + 2\gamma_{xz}\delta S_{xz} + 2\gamma_{yz}\delta S_{yz} + 4\lambda_x \delta l_x + 4\lambda_y \delta l_y + 4\lambda_z \delta l_z - (P-P_0)\delta V,$$

where $\delta S_{ij}$ is the variation of the area of the parallelepiped face lying on the $i-j$ plane, $\delta l_i$ is the variation of the $i-$th side, $P$ is the internal pressure and $P_0$ is the external pressure. In the following we shall truncate fluctuations at the linear order. By differentiating the variation of mechanical work with respect to the variation of length of each of the sides of the parallelepiped and renormalising by the surface normal to that direction, we obtain the normal force densities:

(4)
$$\begin{cases} f_x = -\frac{1}{l_y l_z}\frac{\partial \delta W}{\partial \delta l_x} = (P-P_0) - \frac{2\gamma_{xy,x}}{l_z} - \frac{2\gamma_{xz}}{l_y} - \frac{4\lambda_x}{l_y l_z} \\ f_y = -\frac{1}{l_x l_z}\frac{\partial \delta W}{\partial l \delta_y} = (P-P_0) - \frac{2\gamma_{xy,y}}{l_z} - \frac{2\gamma_{yz}}{l_x} - \frac{4\lambda_y}{l_x l_z} \\ f_z = -\frac{1}{l_x l_y}\frac{\partial \delta W}{\partial \delta l_z} = (P-P_0) - \frac{2\gamma_{xz}}{l_y} - \frac{2\gamma_{yz}}{l_x} - \frac{4\lambda_z}{l_y l_x} \end{cases}.$$

The three non-zero components of the stress tensor are finally given by:

(5)
$$\begin{cases} \sigma_{xx} = 2\eta \frac{1}{l_x}\frac{dl_x}{dt} - (P-P_0) + \frac{2\gamma_{xy,x}}{l_z} + \frac{2\gamma_{xz}}{l_y} + \frac{4\lambda_x}{l_y l_z} \\ \sigma_{yy} = 2\eta \frac{1}{l_y}\frac{dl_y}{dt} - (P-P_0) + \frac{2\gamma_{xy,y}}{l_z} + \frac{2\gamma_{yz}}{l_x} + \frac{4\lambda_y}{l_x l_z} \\ \sigma_{zz} = 2\eta \frac{1}{l_z}\frac{dl_z}{dt} - (P-P_0) + \frac{2\gamma_{xz}}{l_y} + \frac{2\gamma_{yz}}{l_x} + \frac{4\lambda_z}{l_y l_x} \end{cases}.$$

To take into account the muscle action on the cell, we introduce an external drag force, $F_{ext}$. Normally, this force is oriented with an angle $\theta = 6°$ with respect to the $x-$direction (see Main Text and [4]). Yet, for the sake of simplicity, we impose $\theta \simeq 0$ and therefore this force density affects force balance along the $x-$direction only, i.e., along the Anterior-Posterior axis of the worm. Since we observe no relative velocity between the cells of interest and the surrounding tissues, we neglect the friction with the surroundings. The final force balance reads:

(6)
$$\begin{cases} \sigma_{xx} = -\frac{F_{ext}}{l_y l_z} \\ \sigma_{yy} = 0 \\ \sigma_{zz} = 0 \end{cases}.$$

To obtain the equation of motion for the three sides of the paralellepiped, in the presence of muscle contractions, we impose a constant drag speed $v = \frac{L_{AP}(t_f) - L_{AP}(t_0)}{t_f - t_0}$ as in the experimental observations and solve the aforementioned system along with the constraint of volume conservation to obtain $\frac{dl_y}{dt}$, $\frac{dl_z}{dt}$, the external force $F_{ext}$ and the difference between internal and external pressure $P - P_0$. The system is therefore fully determined, with the external force and the pressure difference acting as two Lagrange multipliers respectively ensuring the observed drag speed and volume conservation. In the absence of external forces, the same procedure applies with the difference that instead of imposing a constant drag speed along the Anterior-Posterior axis, we shall impose the external force to be zero and solve the system for $\frac{dl_x}{dt}$ as well.

To take into account the action of cadherins on mechanics, we assume that the concentration of cadherins alters adhesion at a given junction and therefore the line tension [5, 6]. In particular, we assume that:

$$\begin{cases} \lambda_x = \lambda_{AP} = \lambda_0 - \epsilon\phi_{AP}(t)E_b \\ \lambda_y = \lambda_{DV} = \lambda_0 - \epsilon\phi_{DV}(t)E_b \end{cases} \tag{7}$$

where $\lambda_0$ is the basal line tension in the absence of cell adhesion, $E_b$ is the E-cadherin binding energy per unit of molecule, $\phi_{AP}(t)$ (resp. $\phi_{DV}(t)$) is the fluorescence intensity E-cadherin profile on the Anterior-Posterior (resp. the Dorso-Ventral) junctions obtained from FRAP experiments and $\epsilon$ is a proportionality constant to match the theoretical concentrations and experimental fluorescence data. The last line tension, $\lambda_z$, is assumed to be equal to the basal line tension $\lambda_0$ since no E-cadherins are observed on that direction. All surface tensions instead are assumed to be equal to the basal surface tension $\gamma_0$ other than $\gamma_{xy,y} = \mu\gamma_0$, where $\mu$ is a scaling factor, to take into account the actomyosin remodelling observed in [3].

The final system of equations in the presence of muscle contractions is given by:

$$\begin{cases} \frac{dl_y}{dt} = -\frac{\gamma_0}{2\eta V_0}\left(\mu l_y^2 l_x + \frac{v\eta}{\gamma_0} l_y^2 l_z - V_0 + 2\frac{\lambda_y}{\gamma_0} l_y^2 - 2\frac{\lambda_z}{\gamma_0} l_y l_z\right) \\ \frac{dl_z}{dt} = -\frac{\gamma_0}{2\eta V_0}\left(l_z^2 l_x + \frac{v\eta}{\gamma_0} l_z^2 l_y - \mu V_0 + 2\frac{\lambda_z}{\gamma_0} l_z^2 - 2\frac{\lambda_y}{\gamma_0} l_y l_z\right) \\ P = P_0 + \gamma_0\left(\frac{\mu}{l_z} + \frac{1}{l_y} + \frac{2}{l_x}\right) - \frac{v\eta}{l_x} + \frac{2\lambda_y}{l_x l_z} + \frac{2\lambda_z}{l_x l_y} \\ F_{ext} = -(2-\mu)\gamma_0 l_y - \gamma_0 l_z - 3v\eta\frac{l_y l_z}{l_x} + 2\gamma_0\frac{l_y l_z}{l_x} - 4\lambda_x + 2\lambda_y\frac{l_y}{l_x} + 2\lambda_z\frac{l_z}{l_x} \end{cases} \tag{8}$$

where $v$ is the drag speed imposed by muscle contractions.

In the absence of muscle contraction we obtain:

$$\begin{cases} \frac{dl_x}{dt} = -\frac{\gamma_0}{3\eta V_0}\left[(2-\mu)l_x^2 l_y + l_x^2 l_z - 2V_0 + \frac{4\lambda_x}{\gamma_0} l_x^2 - \frac{2\lambda_y}{\gamma_0} l_x l_y - \frac{2\lambda_z}{\gamma_0} l_x l_z\right] \\ \frac{dl_y}{dt} = -\frac{\gamma_0}{3\eta V_0}\left[(2\mu-1)l_y^2 l_x + l_y^2 l_z - 2V_0 + \frac{4\lambda_y}{\gamma_0} l_y^2 - \frac{2\lambda_x}{\gamma_0} l_y l_x - \frac{2\lambda_z}{\gamma_0} l_y l_z\right] \\ \frac{dl_z}{dt} = -\frac{\gamma_0}{3\eta V_0}\left[l_z^2 l_y + l_z^2 l_x - (1+\mu)V_0 + \frac{4\lambda_z}{\gamma_0} l_z^2 - \frac{2\lambda_y}{\gamma_0} l_z l_y - \frac{2\lambda_x}{\gamma_0} l_z l_x\right] \\ P = P_0 + \frac{4}{3}\gamma_0\left[\frac{1+\mu}{2l_z} + \frac{1}{l_y} + \frac{1}{l_x}\right] + \frac{4}{3V_0}\left[\lambda_x l_x + \lambda_y l_y + \lambda_z l_z\right]. \end{cases} \tag{9}$$

This system shows that: i) all mechanical parameters can be rescaled with respect to the viscosity thus reducing to 5 free parameters; ii) the main elongation or contraction speeds are controlled by the ratio between the surface tension and the viscosity [2]; iii) the contribution of the Anterior-Posterior distribution of cadherins in general favours elongation along the Anterior-Posterior axis and thinning of the embryo, whereas the Dorso-Ventral distribution of cadherin favours elongation along the Dorso-Ventral axis and still thinning.

These results are globally reported in the phase diagrams shown Fig. S6A-C; iv) the presence of an anisotropy in surface tension leads to a constriction along the Anterior-Posterior and Dorso-Ventral axis and thickening of the embryo (Fig. S6D).

#### 1. Numerical simulations and parameter estimates

We solved these systems of equations by implementing a numerical scheme in Matlab [7] using the ordinary differential equation solver `ode15s()` for stiff systems. To this aim, we first solved the cadherin ordinary differential equations and then fed the mechanical systems with these profiles to obtain the final solutions for the embryo deformations. We used this algorithm to first reconstruct the phase diagrams of deformations as a function of the ratio of the Anterior-Posterior and Dorso-Ventral kinetic parameters for the E-cadherins as well as of the myosin cable contribution to the surface tension, $\mu$ (Fig. S6A-D).

In order to test the model ability to quantitatively reproduce Wild Type experiments in the body and head cells respectively, we fixed the cadherin kinetics as obtained from FRAP experiments and the cadherin binding energy to $E_b = k_B T$[5], where $k_B$ is the Boltzmann constant and $T$ is the room temperature. We imposed the elongation dynamics along the Anterior-Posterior axis by fixing the drag speed $v$ following the quantifications of the Anterior-Posterior length over time respectively obtained in the body and in the head. We assumed that the initial conditions for each experiment were given by the data, other than $l_z(0)$ assumed to be $l_z(0) = 2$ $\mu$m. We implemented a search algorithm in Matlab based on the `fmincon()` function [7] aimed at minimising the Euclidean distance between the outcome of the simulations and the data quantifying the Dorso-Ventral deformations as a function of the mechanical parameters ($\gamma_0/\eta$,$\lambda_0/\eta$ and $\mu$) and the proportionality constant between cadherin concentration and fluorescence $\epsilon/\eta$. The algorithm was constrained such that the line tensions be always positive as a function of the different mechano-chemical parameters. We started by applying this algorithm to the Wild Type body data. We repeated the minimisation $N_{sample} = 50$ times and extracted the mean and the coefficients of variation (standard deviation-to-mean ratio) of the obtained solutions for each parameter, contained in the vector $\pi^*$=($\gamma_0^*/\eta$,$\lambda_0^*/\eta$ and $\mu^*$). The results are reported in Table S1 and Fig. S6I-J and they show that the algorithm robustly inferred these parameters, all of the coefficients of variation being less than 1. To obtain the same estimates for the Wild Type head data, we imposed the same parameter $\epsilon$ as in the Wild Type body data and simply assumed a rescaling constant $\kappa$ for all the remaining mechanical parameters: this allowed us to now perform the same search by minimising the same objective function with respect to one parameter only. The result is reported in Table S1 and Fig. S6I-J and shows that, as expected from previous measurements, the mechanical parameters in the Wild Type head are higher than those in the Wild Type body [3].

To test the predictiveness of the model, we then used the parameters obtained in the previous experiments on Wild Type and the quantifications of the cadherin kinetics from FRAP experiments for the Pat mutants to run the simulations in the absence of external force. As shown in Fig. 6B-I, the simulations well reproduced the mutant quantifications in the body and sufficiently well the evolution of the Dorso-Ventral side in the head, thus confirming the predictiveness of the model and of its parameter inference.

The final full set of parameters is reported in Table S1 and discussed in the Main Text.

#### 2. Effect of E-cadherin kinetics alteration on elongation

In order to understand how alterations in E-cadherin kinetics, i.e., how the values of the endo- and exocytosis rates affect eventually cell mechanics and therefore its elongation, we ran simulations according to the following protocol.

We rescaled by a factor $\sigma = 0.1, 0.2, 0.4$ and $0.8$ either the endocytosis rate $\mathrm{k}_{\mathrm{off}}$ or the exocytosis rate $\mathrm{k}_{\mathrm{on}}$ along both the Anterior-Posterior and the Dorso-Ventral axis. With these new values of E-cadherin exocytosis and endocytosis rate, we first numerically integrated Eqs. 1-3 of the Main Text to obtain the fluorescence time profile . We then used these profiles as an input for the numerical integration of the mechanical equations Eqs. 8 of this Supplementary Theory. All parameters are the same as in WT body simulations other than E-cadherin endo- and exocytosis rates and the basal line tension $\lambda_0$, which has been multiplied by 4 in order to prevent the total line tensions to become negative. The results are reported in Fig. S6L-O and commented in the Main Text.

[7] "The mathworks inc., matlab version: 23.2.0.2485118 (r2023b)," 2023.

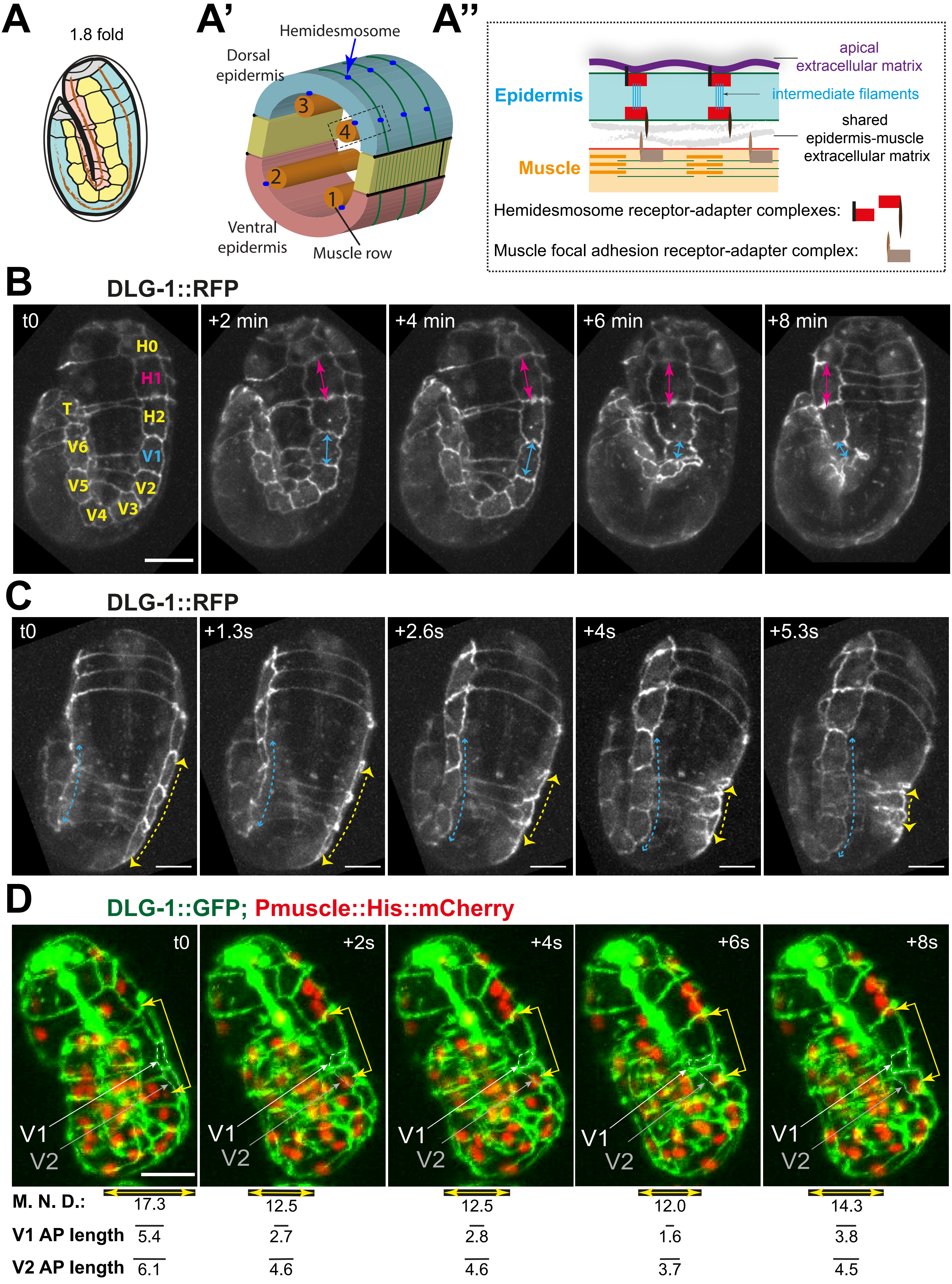


### Supplementary Figure 1

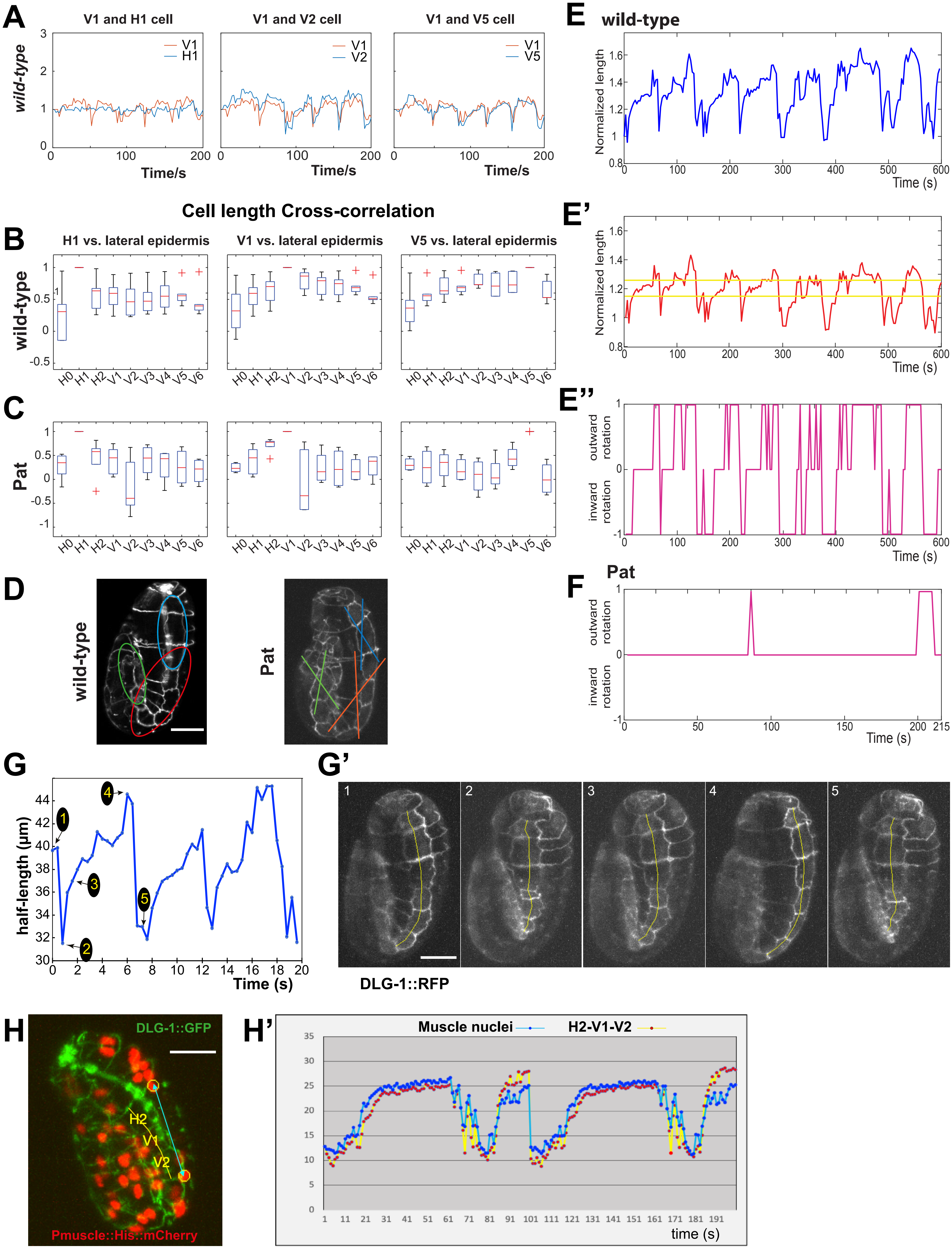


### Supplementary Figure 2

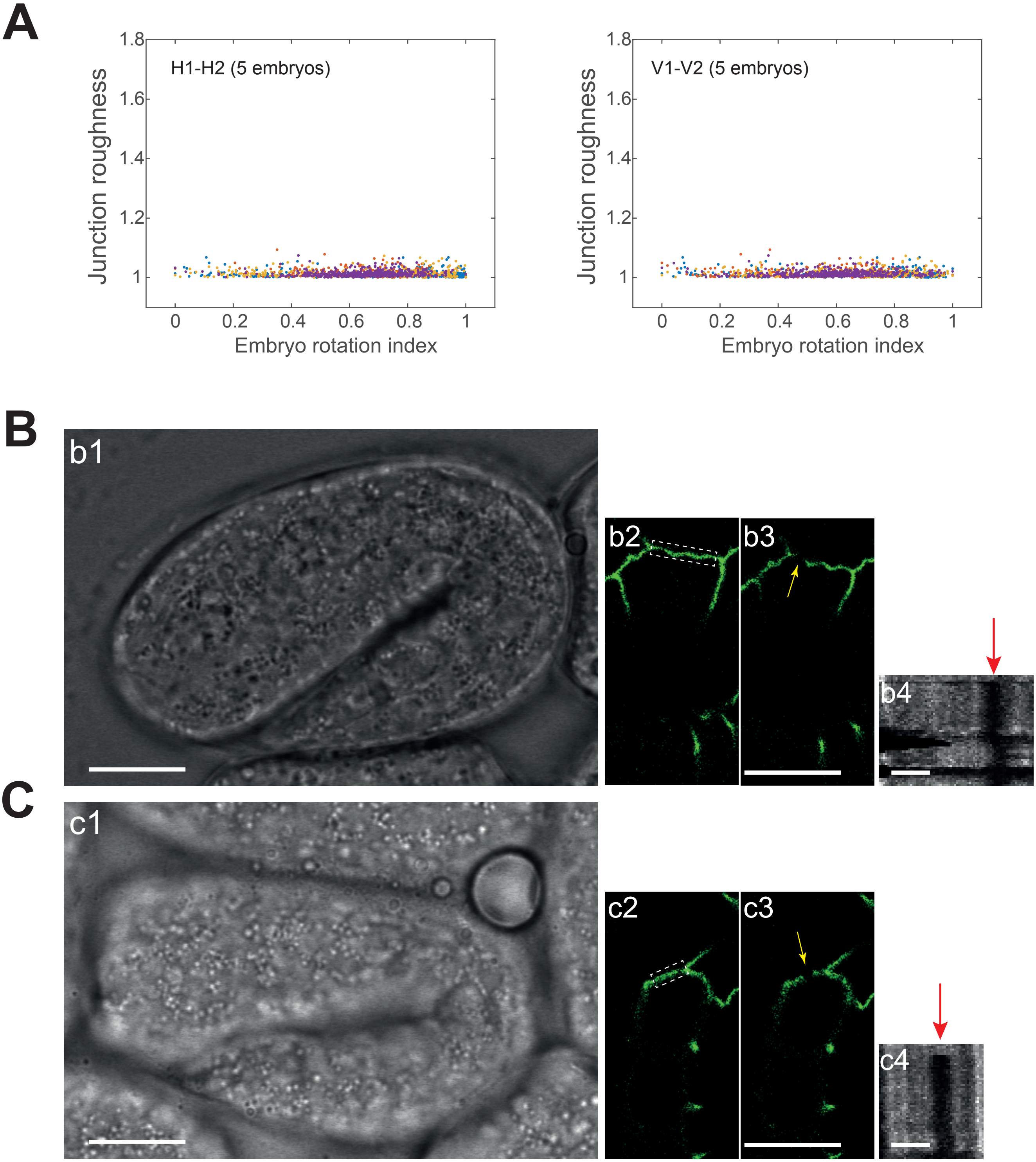


### Supplementary Figure 3

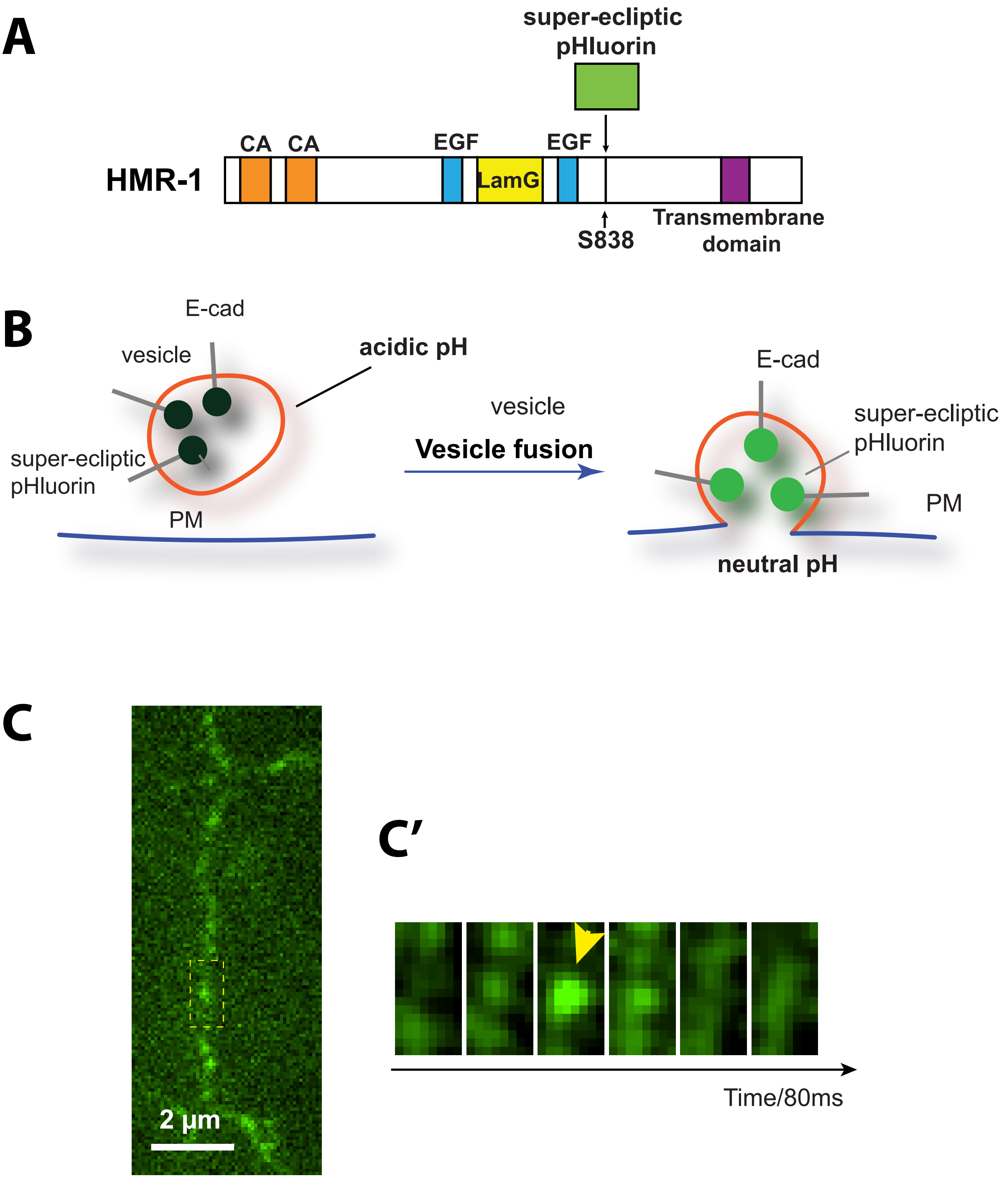


### Supplementary Figure 4

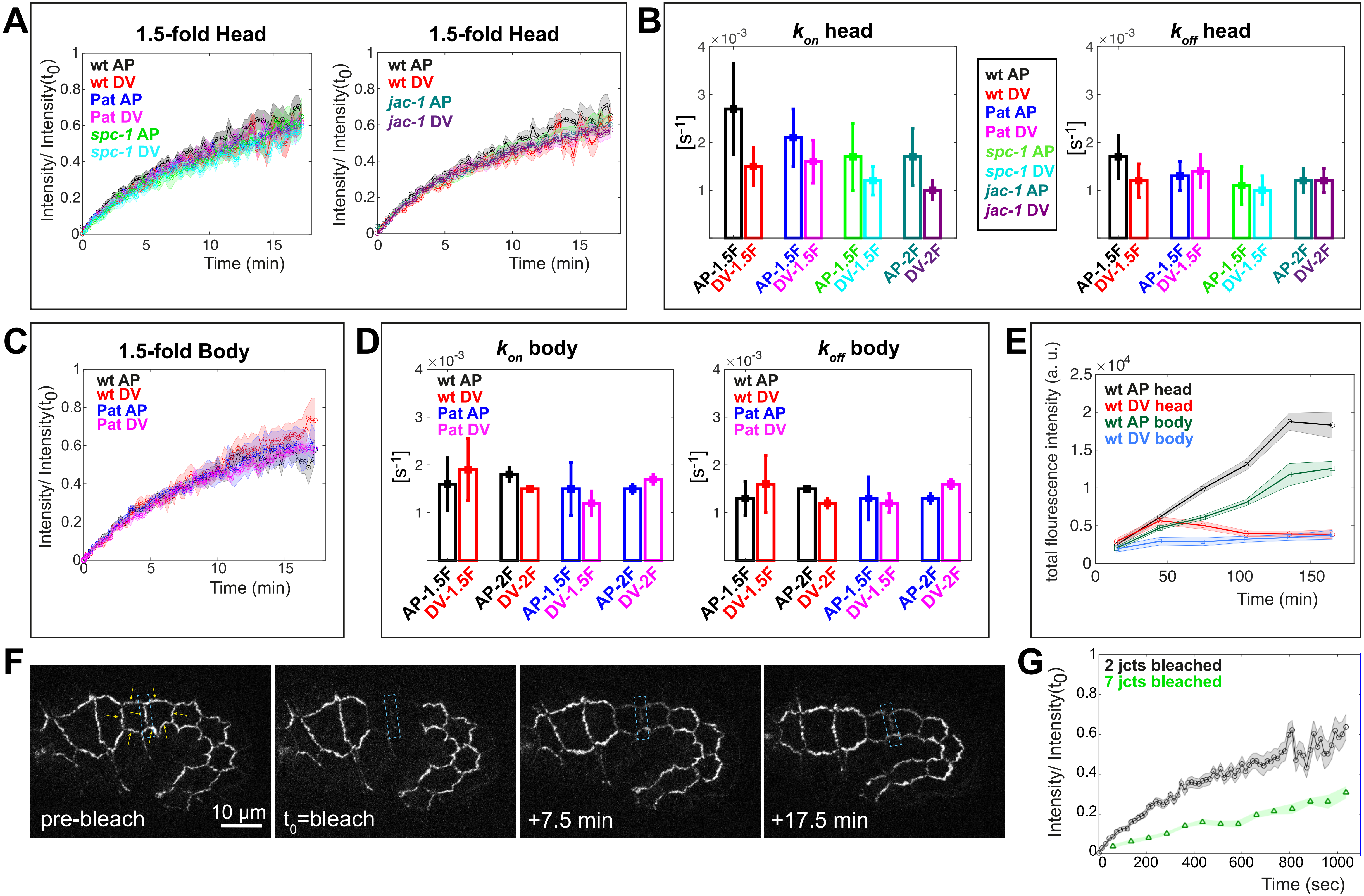


Supplementary Figure 5

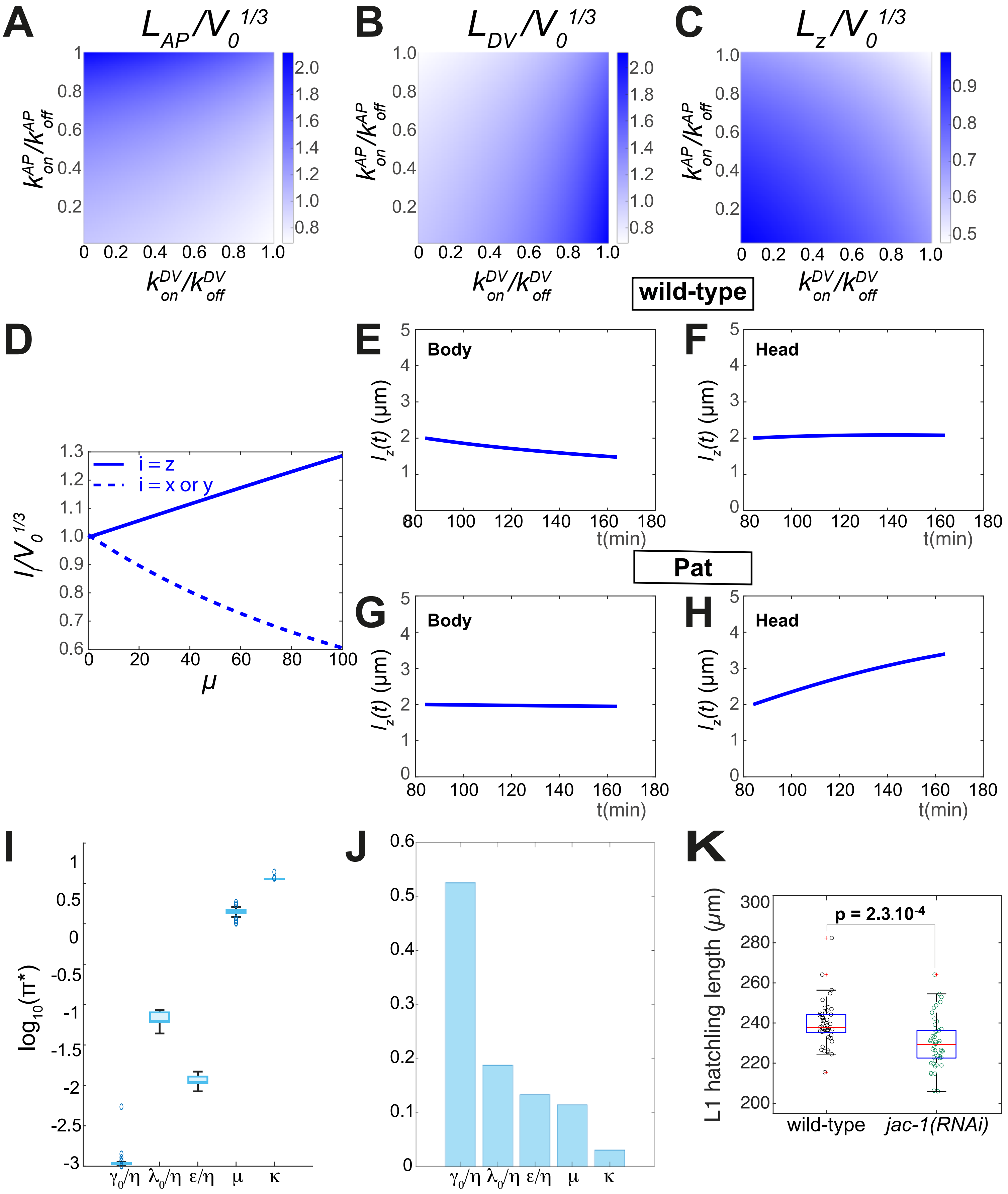


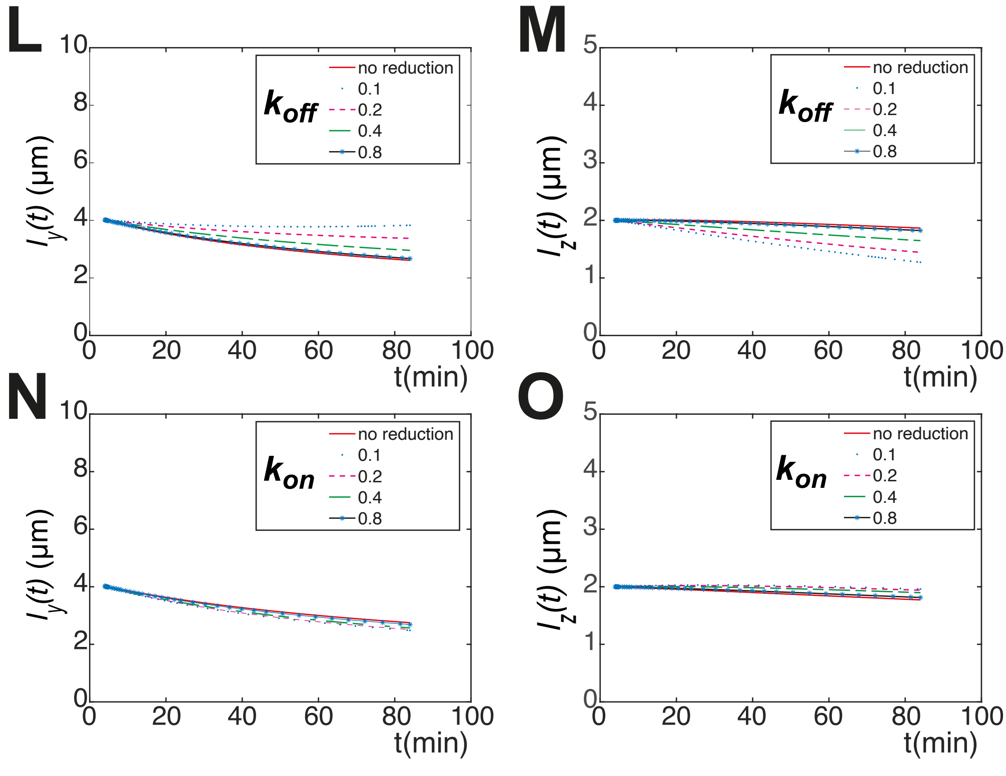


P

| | WT body | WT head |
|---|---|---|
| Muscle drag force/ line tension **AP** jct = $\langle Fext \rangle_t / (\varepsilon \langle \phi_{AP} \rangle_t Eb)$ | 2.46 | 2.45 |
| Muscle drag force/ line tension **DV** jct = $\langle Fext \rangle / (\varepsilon \langle \phi_{DV} \rangle_t Eb)$ | 13.97 | 17.80 |

### Supplementary Figure 6
